# The bistable mitotic switch in fission yeast

**DOI:** 10.1101/2024.01.23.576917

**Authors:** Béla Novák, John J. Tyson

## Abstract

In most situations, eukaryotic cells proceed irreversibly through the cell division cycle (G1-S-G2-M) in order to produce two daughter cells with the same number and identity of chromosomes of their progenitor. The integrity of this process is maintained by ‘checkpoints’ that hold a cell at particular transition points of the cycle until all requisite events are completed. The crucial functions of these checkpoints seem to depend on irreversible bistability of the underlying checkpoint control systems. Bistability of cell cycle transitions has been confirmed experimentally in frog egg extracts, in budding yeast cells and in mammalian cells. For fission yeast cells, a recent paper by Patterson et al. (1) provides experimental evidence for an abrupt transition from G2 phase into mitosis, and we show that these data are consistent with a stochastic model of a bistable switch governing the G2/M checkpoint. Interestingly, our model suggests that their experimental data could also be explained by a reversible/sigmoidal switch, and stochastic simulations confirm this supposition. We propose a simple modification of their experimental protocol that could provide convincing evidence for (or against) bistability of the G2/M transition in fission yeast.

## Introduction

Proliferating eukaryotic cells alternate between two phases: interphase and mitosis (M phase). Throughout interphase (G1-, S-and G2 phases) chromosomes are decondensed. During S phase every chromosome is replicated to produce two identical sister chromatids held together by cohesin rings. In contrast, chromosomes are condensed during M phase, which prepares sister chromatids for segregation during anaphase (2). Maintaining the ploidy of the cell requires that chromosome-replicating interphase strictly alternates with chromatid-segregating M phase. This requirement is satisfied by the irreversibility of the transitions between interphase and M phase (at G2/M boundary) and mitotic exit (anaphase-telophase-cell division). In 1993, we proposed that irreversibility of these transitions is a consequence of bistability in the molecular mechanism controlling entry into and exit from mitosis (3, 4). This mechanism is based on cyclin activation of mitosis promoting factor (MPF). The accumulation of ‘mitotic’ cyclin in interphase triggers abrupt activation of MPF and entry into mitosis (5). Degradation of mitotic cyclin in anaphase and telophase leads to an abrupt loss of MPF activity as the cell divides and enters G1 phase. ‘Bistability’ of the mitotic switch means that the cyclin threshold for mitotic entry is larger than the threshold for mitotic exit; for cyclin levels between these two thresholds, the M-phase control system can be either in an interphase state (low level of MPF activity) or in a mitotic state (high level of MPF activity). For an interphase cell to enter mitosis, the level of mitotic cyclin must increase above the ‘entry’ threshold, and then, to return to interphase, mitotic cyclin must be degraded to a level below the ‘exit’ threshold. We proposed that this behavior, known as hysteresis, is the basis of the alternation between interphase and mitosis during cell proliferation (3).

Based on the evolutionary conservation of the mitotic control network (6), we applied this concept of a bistable mitotic switch to both the free-running cell cycles of early Xenopus embryos (non-growing cells) and to the growth-controlled cell cycles of fission yeast (3, 4, 7). Dedicated experiments with Xenopus cell-free extracts confirmed these predictions by showing that more mitotic cyclin is required for mitotic entry than for maintaining the extract in M phase (8, 9). Later, we applied the bistability concept to the G1/S transition in budding yeast and mammalian cells (10–12). The distinctive signatures of bistability were observed experimentally in budding yeast (13, 14) and in a human (HeLa) cell line (15, 16), suggesting that bistability is a conserved feature of the mitotic switch.

Interestingly, experimental evidence for bistability of mitotic control in fission yeast has been missing. In 2021, Nurse’s group measured cyclin-CDK <u>activity</u> (i.e., MPF) as a function of total cyclin-CDK <u>level</u> in fission yeast cells and observed that cells close to the G2/M transition (cell length ≈ 12 μm) and containing equivalent amounts of cyclin-CDK could be either in G2 phase (low cyclin-CDK activity) or in M phase (high cyclin-CDK activity), which they interpreted as evidence that the dose-response curve is “clearly bistable, with cells existing in either an ‘on’ or an ‘off’ state” (1). Intrigued by these results, we investigated their experimental observations with a model of bistability in the activation of cyclin-CDK in fission yeast. Using a reliable stochastic simulation algorithm (17), we show that our model yields dose-response curves in near-perfect agreement with Patterson et al.’s observations of ‘wildtype’ cells and three mutant strains with aberrant activation of cyclin-CDK. Our analysis of these simulations and the experimental data shows, furthermore, that Patterson’s protocol probes only the cyclin threshold for mitotic entry (i.e., CDK activation) and not the cyclin threshold for mitotic exit (CDK inactivation). Therefore, the experiments do not provide unequivocal evidence for bistability. Indeed, we show that Patterson et al.’s experimental data are also consistent with a reversible (ultrasensitive, not bistable) mitotic transition, and we propose a modified experimental protocol that could provide unambiguous evidence for bistability in the mitotic control system of fission yeast.

### The experiment

To study the dependence of mitotic entry on the accumulation of mitotic CDK activity in fission yeast, Patterson et al. (1) created a strain carrying a temperature-sensitive Cdk1 mutation (*cdc2^ts^*) and a tetracycline-inducible, GFP-tagged Cdc13^dbΔ^-Cdk1 fusion protein (*TetP*:*cdc13^dbΔ^*-*GFP*-*cdc2*). The cyclin component of the fusion protein lacked the destruction box of the wildtype *cdc13^+^* gene; hence, their fusion protein is not degraded after the induced cell enters mitosis. Cells were also equipped with a fluorescently-tagged Cut3-based CDK activity sensor (*synCut3-mCherry*), which accumulates in the nucleus in its CDK-phosphorylated form. In their experiments, Patterson et al. first raised cells to the restrictive temperature to inactivate the cell’s endogenous supply of wildtype Cdk1 and then induced synthesis of the Cdc13^dbΔ^-Cdk1 fusion protein (denoted C-CDK, with the understanding that the cyclin component is non-degradable; a distinction that will become important later). Using single-cell assays, they simultaneously measured C-CDK level (by GFP), CDK activity (by mCherry) and cell size at increasing times after induction of the fusion protein. The level of induced C-CDK in a cell depended on the length of induction rather than cell size (as in normal cell-cycle progression). In this way Patterson et al. managed to characterize the dependence of the mitotic cyclin threshold (i.e., fusion protein level) for CDK activation on cell size. As a check on the underlying mechanism of the mitotic control system, they also studied a form of the fusion protein, Cdc13^dbΔ^-Cdk1^AF^ (denoted C-CDK^AF^), that cannot be inactivated by inhibitory phosphorylation of the Cdk1 subunit by Wee1/Mik1 kinase. In addition, they did experiments in a *ppa2Δ* genetic background, where one of the type-2A protein phosphatases (PP2A) has been deleted. In their ‘wildtype’ strain (i.e., *cdc13^dbΔ^-cdc2^+^ppa2^+^*), the authors observed a wide range of fusion protein levels where cells with both low and high CDK activity coexist.

Patterson et al. (1) interpreted their observations as in vivo confirmation of the bistable “C-CDK activity vs C-CDK level dose-response curves previously demonstrated in vitro” by Sha et al. (8) and Pomerening et al. (9) in frog egg extracts. Although we agree with Patterson et al. that the mitotic control system in fission yeast is (as we predicted) governed by a bistable switch, we will show here that their data corresponds more closely to an early study in frog egg extracts by Solomon et al. (5), who observed a sigmoidal dependence of C-CDK (‘MPF’) activity on C-CDK level (‘total cyclin’). Neither the experiments of Solomon et al. in vitro nor those of Patterson et al. in vivo provide proof of ‘bistability’ because these experiments probe only one half of the dose-response curve of a bistable switch (turning on the switch by raising the cyclin level). The experiments of Sha et al. and Pomerening et al., based on our earlier computational studies (3), provide convincing evidence of bistability of the mitotic switch by raising and lowering the cyclin level in frog egg extracts and showing that, at an intermediate level of cyclin, the extract could remain in a stable steady state of low or high MPF activity depending on whether the extract began in a state of low or high MPF activity (a phenomenon called ‘hysteresis’). By using stochastic and deterministic modelling of the fission-yeast mitotic switch, we show that the observations of Patterson et al. (1) are consistent with both an irreversible/bistable mitotic switch and a reversible/sigmoidal mitotic switch. Although we are quite sure the bistable switch is the correct interpretation, a sigmoidal switch is not excluded by the experiments. To resolve this uncertainty, we propose a simple modification to Patterson’s protocol that would allow the C-CDK level to both rise and fall and thereby provide unequivocal evidence for or against a bistable mitotic switch.

## Results and Discussion

### The model

Our model of the mitotic control system in fission yeast is based on our previous work (18–21). C-CDK is phosphorylated on inhibitory sites of Cdk1 by Wee1/Mik1 kinases, and these phosphorylations are reversed by active Cdc25-phosphatase (Fig.1). Both the kinases and the phosphatase are phosphorylated by active C-CDK, but with opposite effects: Wee1 phosphorylations are inhibitory, Cdc25 phosphorylations are activatory. C-CDK phosphorylation of these regulatory enzymes are reversed by dephosphorylation by PP2A (18). In our dynamic description of Wee1/Mik1-catalyzed phosphorylation of C-CDK we explicitly include the complex, Wee1:CDK, formed by the inhibitory kinases and their substrate (see Fig.1). This novel feature of our model means that Wee1 and Mik1 function as stoichiometric inhibitors of C-CDK even when the Cdk1 subunit cannot be phosphorylated (i.e., the C-CDK^AF^ strain). Consistent with experimental observations, our model has Cdc25 level increasing with cell size (22, 23). Also, the level of Wee1/Mik1 is cell cycle regulated, for the following reason: although Wee1 is maintained at constant level, Mik1 protein level peaks early in the cycle due to induction by a cell cycle-regulated transcription factor, MBF (24, 25).

**Figure 1:**
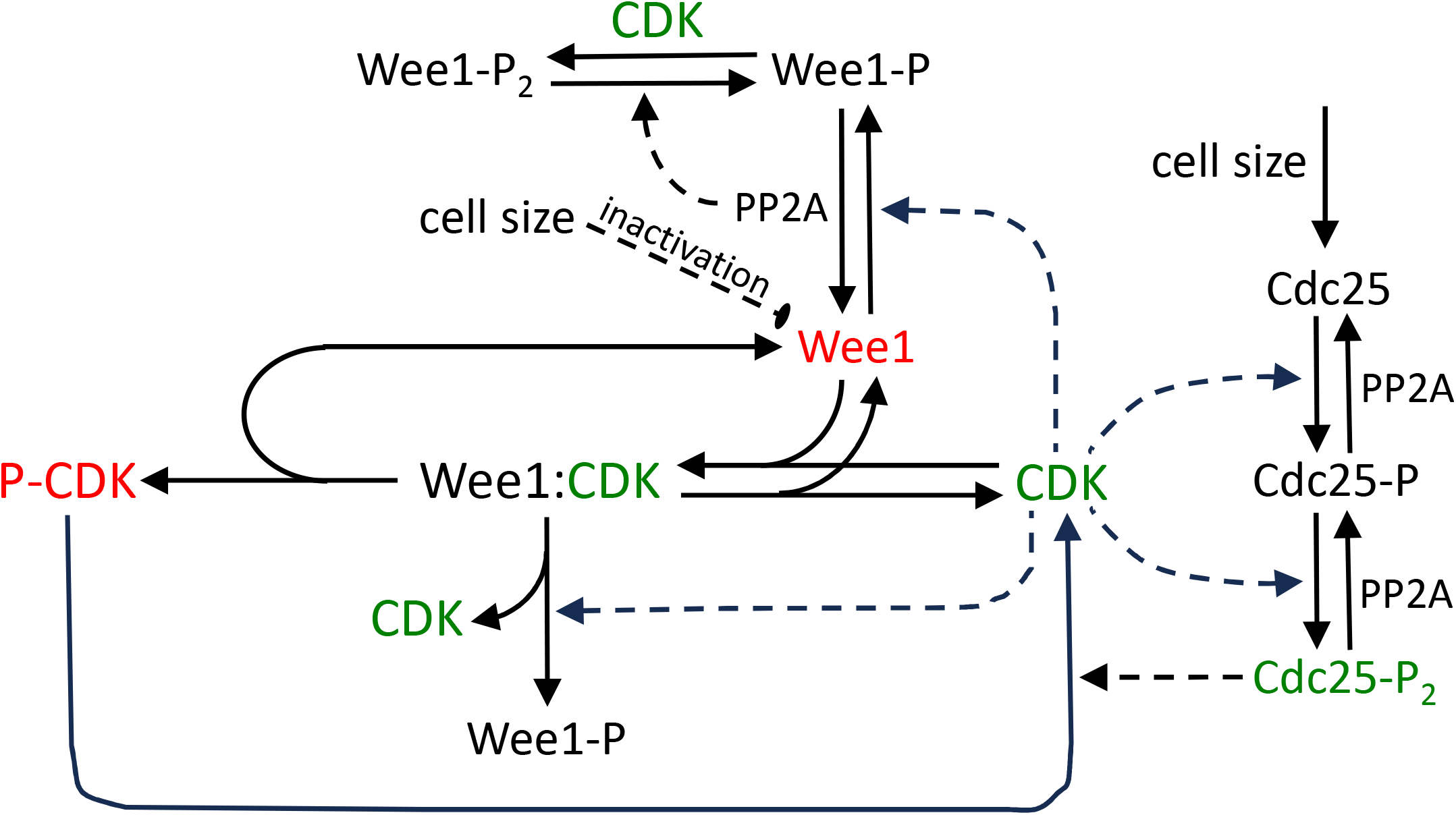
The molecular regulatory network of the model. CDK is inhibited by phosphorylation by Wee1/Mik1 kinases and reactivated by Cdc25 phosphatase. Wee1/Mik1 kinases form a stoichiometric complex with CDK before catalyzing its inhibitory phosphorylation. Both Wee1/Mik1 and Cdc25 are doubly phosphorylated by CDK, and the phosphorylations are reversed by PP2A. Because the active forms of Wee1/Mik1 are unphosphorylated and the active form of Cdc25 is doubly phosphorylated, the net activation of CDK is governed by two interlocked, positive feedback loops.

The model was also supplemented with a variable that accounts for the CDK activity sensor. Guided by experimental data, we assume that the unphosphorylated sensor rapidly equilibrates between cytoplasm and nucleus, but nuclear export of the phosphorylated form is attenuated (see Materials and Methods).

### Stochastic simulations

We simulated the temporal dynamics of the network in Fig.1 with Gillespie’s Stochastic Simulation Algorithm (SSA) (17). Because our model of molecular interactions is formulated in terms of mass-action rate laws, Gillespie’s SSA provides reliable simulations of the stochastic fluctuation of the biochemical control system. Simulations were run with different initial cell volumes for different lengths of time, allowing synthesis of fusion protein (C-CDK) during exponential volume growth. In the model, the rate of synthesis of C-CDK molecules is directly proportional to cell volume, and the molecules have a long half-life, because the fusion protein has a deleted destruction box. Data for cell volume, C-CDK concentration and nucleocytoplasmic ratio of the CDK-activity sensor were collected at each time point. In this way stochastic simulations of the model provided predictions of how CDK activity depends on cell size and fusion protein levels in single cells, as observed in the experiments. Each genetic background was analyzed by more than 6,000 simulations.

The nucleocytoplasmic ratio of the CDK-activity sensor, plotted as a function of C-CDK concentration in our Fig.2, is directly comparable with the original experimental data in Fig.2i of Patterson et al. (1). We have grouped the cells into four cell-size bins (V=0.5-0.65, V=0.65-0.8, V=0.8-0.95 and V=0.95-1.1) similar to the presentation of experimental results in Patterson et al. (1). Our simulations of phosphorylable and non-phosphorylable C-CDKs are provided on the left and the right panels of Fig.2, whereas the top and the bottom panels show the cases of *pp2a^+^* and PP2A-deleted cells. In Suppl. Fig.S1, we plot the same simulations in terms of the level of the active form of C-CDK. While the CDK activity measured by the sensor saturates at high fusion-protein levels, consistent with experimental data, the level of active CDK increases linearly in the same region.

**Figure 2:**
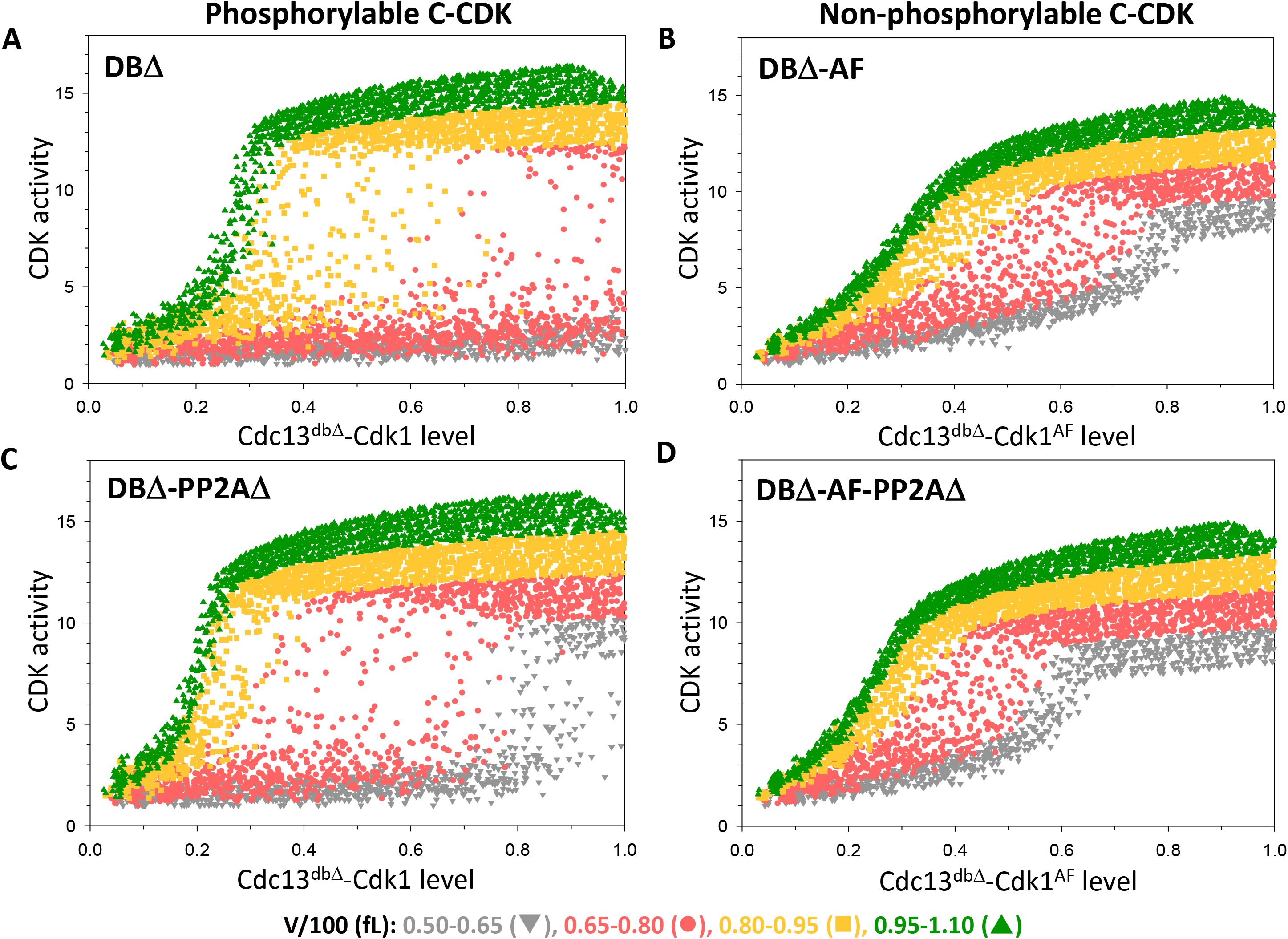
Stochastic simulations of the bistable mitotic-switch model. CDK activity is plotted as a function of fusion-protein level after induction of Cdc13^dbΔ^-Cdk1 (left column: A & C) and Cdc13^dbΔ^-Cdk1^AF^ (right column: B & D) in *pp2a*^+^ (top row: A & B) and *pp2aΔ*-deleted background (bottom row: C & D). Cells are sorted into four cell size bins.

Based on CDK activities, three different types of cells (low, intermediate or high CDK activity) can be distinguished in each situation. Cells with low and high CDK activities are, respectively, before and after the process of CDK activation. At very small and very high C-CDK levels, all cells have, respectively, very low and very high CDK activity regardless of cell size. Cells with intermediate CDK activities are in the process of activating CDK, because no cell inactivates its CDK during simulations (or in the experiments). In both genetic backgrounds (*pp2a^+^* and *ppa2Δ*) and with both forms of fusion protein (phosphorylable and non-phosphorylable), CDK becomes activated at smaller fusion protein levels in large cells (green) than in smaller cells (yellow and red) (Fig.2 and Suppl. Fig.S1), clearly indicating that the cyclin (fusion protein) threshold for CDK activation is cell-size dependent. The lack of inhibitory CDK phosphorylation in the AF strain (CDK^AF^) reduces the cyclin threshold for CDK activation more than deletion of PP2A (Fig.2B,C). The reduction of the size thresholds by CDK^AF^ and PP2AΔ are additive (the AF-PP2AΔ strain in Fig.2D).

### Deterministic model

To gain some insight into the stochastic simulations in Fig.2, we analyze the corresponding deterministic model by means of one-parameter bifurcation diagrams (Fig.3) for each of the four strains (DBΔ, DBΔ-AF, DBΔ-PP2AΔ, and DBΔ-AF-PP2AΔ). Each bifurcation diagram depicts the dependence of the steady state value of the CDK activity sensor on the total concentration of fusion protein in a cell. (The corresponding bifurcation diagrams for the level of the active form of C-CDK are provided in Suppl. Fig.S2.) For each strain, the dependence of CDK activity on fusion protein level is calculated for four cell volumes, corresponding to the middle values of the cell bins (V=0.575, 0.725, 0.875 and 1.025) in the stochastic simulations.

**Figure 3:**
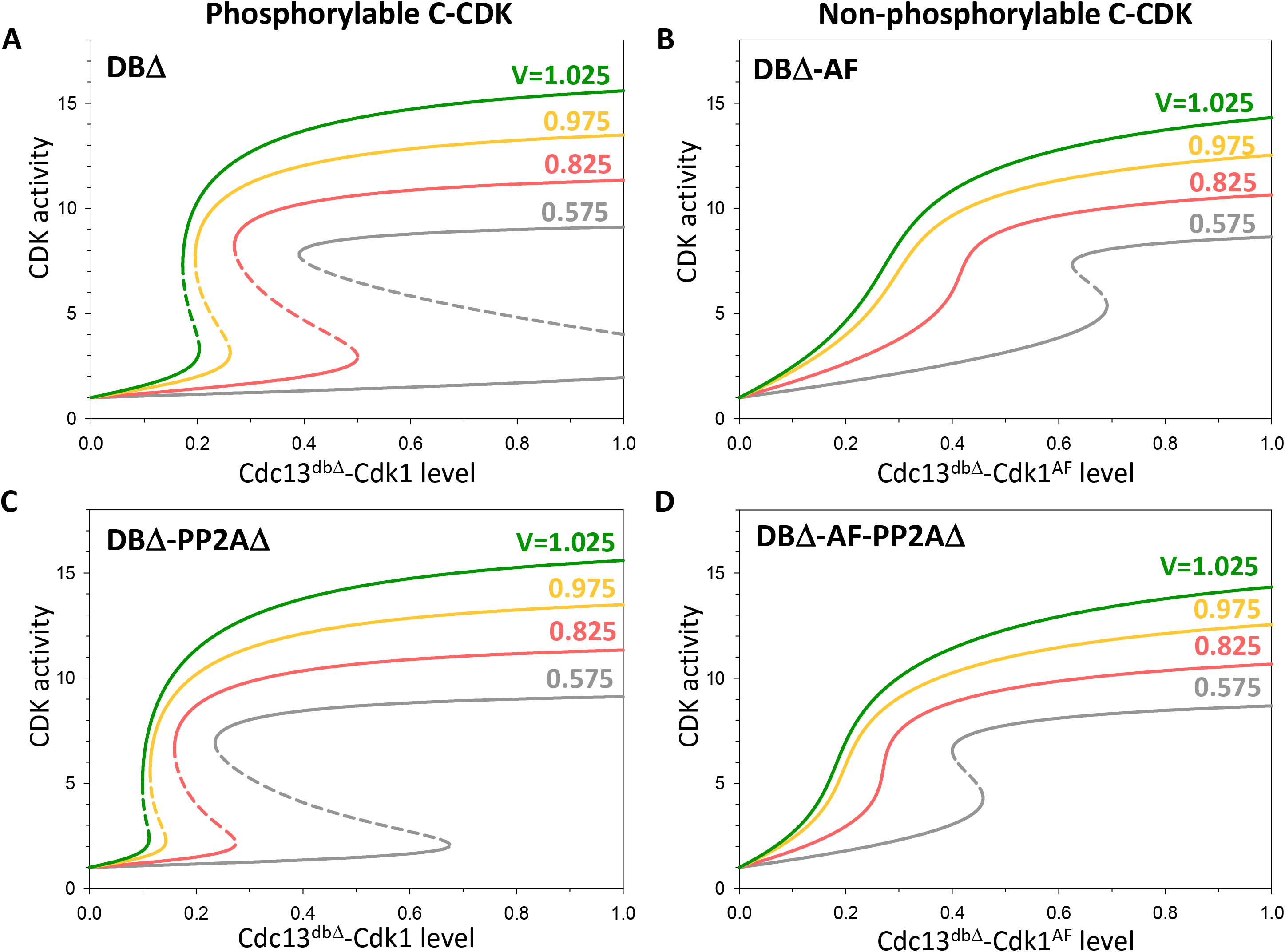
Dose-response curves for the dependence of the CDK-activity sensor on fusion-protein level predicted by the deterministic model. CDK-activity sensor is plotted as a function of fusion-protein level after induction of Cdc13^dbΔ^-Cdk1 (left column: A & C) and Cdc13^dbΔ^-Cdk1^AF^ (right column: B & D) in *pp2a*^+^ (top row: A & B) and *pp2aΔ*-deleted background (bottom row: C & D). CDK activities are plotted for fixed values of V in the middle of the cell-size bins defined for the stochastic simulations.

In the phosphorylable C-CDK strains, the CDK activity vs. fusion protein level is S-shaped at all four sizes; whereas, for the non-phosphorylable C-CDK strains, the bifurcation curves are S-shaped only for the smallest cells. The lower and the upper branches of the S-shaped curves are stable steady states corresponding to interphase and M-phase activity of CDK. In the phosphorylable C-CDK strains, the interphase and mitotic states are overlapping at intermediate levels of fusion protein, indicating bistability in the mitotic control system. The low and high CDK activity states are connected by a middle branch of unstable steady states where the mitotic control system cannot stay for too long in a stochastic environment. The termination of the lower steady state corresponds to the cyclin (fusion protein) threshold for CDK activation; this threshold decreases dramatically as a function of cell volume for the non-phosphorylable C-CDK strains. At the other end of the bistable range, the termination of the high CDK activity state represents the mitotic cyclin threshold for CDK inactivation. In summary, at the heart of our model for the fission yeast mitotic control is a strong bistable switch with two different fusion-protein thresholds between interphase and M-phase transition. Small cells of the strains carrying the Cdk1^AF^ allele exhibit weak bistability in the absence of inhibitory CDK phosphorylation because of mutual antagonism remaining between Cdk1^AF^ and Wee1: unphosphorylated Wee1 binds to and inhibits Cdk1^AF^, and Cdk1^AF^ phosphorylates and inactivates Wee1.

The range of bistability is dependent on cell size as well as genetic background and biochemical details. This dependence is best illustrated on a two-parameter bifurcation diagram, which depicts the boundaries between interphase and mitosis as functions of fusion protein level and cell size (Suppl. Fig.S3). High fusion protein level and large cell size favor the mitotic state, while low fusion protein level and small cell size keep cells in interphase. The cyclin (fusion protein) thresholds for CDK activation and inactivation (the green and the red curves, respectively, in Fig.S3) correspond to the boundaries of interphase and mitotic states, and both thresholds are decreasing functions of cell size. As cell size increases, the two curves meet at a ‘cusp’ point, which represents a merger of interphase and mitotic states at large cell size. The range of bistability can be estimated by the horizontal difference between the two curves, which is illustrated by the dotted line at V=0.6. Figure S3B shows that, in the absence of inhibitory CDK phosphorylation (Cdk1^AF^), bistability is exhibited only by small cells (V<0.7) and only over a narrow range of C-CDK levels. Notice that the absence of PP2A shifts bistability to smaller C-CDK ranges (Fig.S3C) and the two effects (Cdk1^AF^ and PP2AΔ) are additive (Fig.S3D).

In the absence of inhibitory CDK phosphorylation, interphase and M-phase are not separated by a bistable switch with sharp thresholds, except for very small cells. Nonetheless, based on the data of Patterson et al. (1), a boundary between interphase and M-phase can be defined by points where CDK activity=5, which is indicated by the dashed lines in Fig. S3B,D.

To characterize their C-CDK activity data, Patterson et al. (1) proposed an informative statistical measure: “the threshold C-CDK level required for 50% of cells to reach a C-CDK activity determined as being >5 in arbitrary units … within different size bins.” This measure is plotted as a function of cell length for their four different strains in Fig.3B of Patterson et al. (1). Their figure is directly comparable to our model if we swap the axes of the two-parameter bifurcation diagram (Fig.S3 becomes Fig.4), and we plot the C-CDK level required to cross CDK activity=5 as a function of cell size (the blue squares on Fig.4 are the experimental data points in Patterson’s Fig.3B). In the bistable regime (Fig.4A,C), the experimental data lie slightly above the mitotic entry threshold of the bistable switch because of a short time-delay in CDK activation. When cell cycle progression is not controlled by bistability (non-phosphorylable C-CDK^AF^, Fig.4B,D), the experimental points are spread around the CDK activity=5 line calculated by the deterministic model. For our stochastic simulations, we plot— for each of the four cell volume classes—the C-CDK level where 50% of cells are above and 50% below CDK activity=5, and these points (the orange circles on Fig. 4) are in excellent agreement with the experimental data in all genetic backgrounds. We conclude that our bistable model provides a good quantitative fit of the experimental results reported by Patterson et al. (1). But the agreement between experiments and modelling also clearly indicates that experiments only scan the dependence of the mitotic entry threshold (the green curve) on cell size. That there is a lower threshold for mitotic exit (the red curve)—and hence a region of bistability—is not probed by the experimental protocol of Patterson et al. (1).

**Figure 4:**
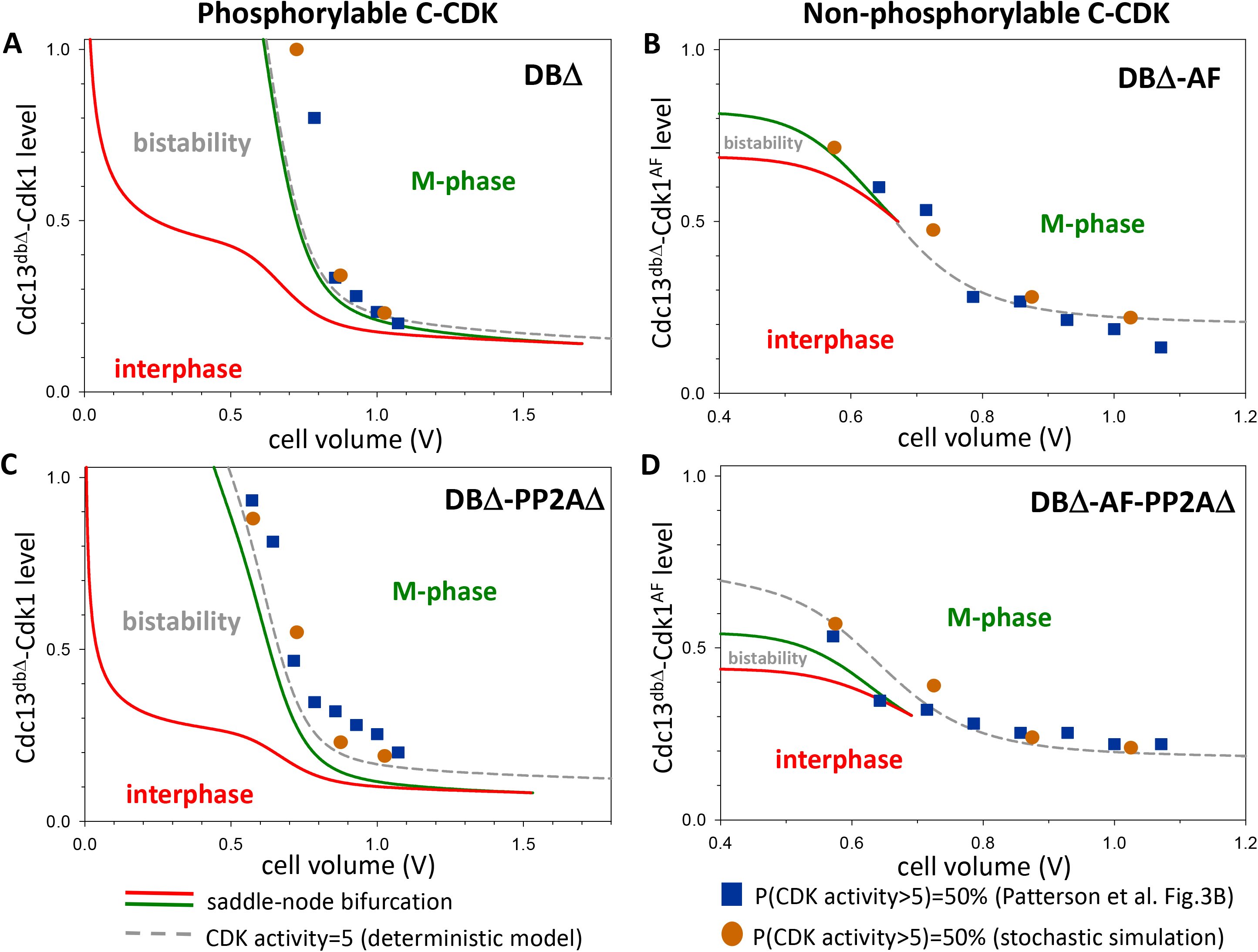
Cell-size dependence of fusion-protein thresholds for mitotic entry and exit in the deterministic model. Induction of Cdc13^dbΔ^-Cdk1 (left column: A & C) and Cdc13^dbΔ^-Cdk1^AF^ (right column: B & D) in *pp2a*^+^ (top row: A & B) and *pp2aΔ*-deleted background (bottom row: C & D). Solid lines depict the saddle-node bifurcation points for transitions into M phase and back to interphase (green and red, respectively). Grey dashed lines depict the fusion protein level where CDK activity reaches the value of 5. Orange circles and blue squares show fusion protein levels where more than 50% of cells have CDK activity higher than 5 in our stochastic simulations and in Patterson et al. (1) experiments, respectively.

### Dynamics of CDK activation in Patterson et al.’s experiments

To understand the dynamics of CDK activation in the experiments of Patterson et al. (1), we analyze the stochastic simulations (Fig.2) in the context of deterministic bifurcation curves (Fig.3). Figure 5A shows a representative stochastic simulation of CDK activation in a growing cell of size V=600 µm^3^ at the start of fusion-protein induction. Since expression of fusion protein starts after inactivation of the endogenous Cdk1 molecules (*cdc2^ts^*), the CDK activity sensor equilibrates between cytoplasm and nucleus and shows an initial value of one (green curve, right axis). Wee1 is predominantly in its unphosphorylated, active form (Fig.5A), which keeps the cell on the lower (interphase) branch of the bifurcation diagram (Fig.5B), with CDK activity near one. Because fusion protein is being synthesized by the cell, the total C-CDK level moves to the right on both diagrams (Fig.5A,B), and this is accompanied by an increase of cell volume, which causes a reduction in the cyclin (fusion protein) threshold for CDK activation (the S-shaped curves in Fig.5B are drifting to the left as the cell grows). When the fusion protein level reaches the right knee of the S-shaped curve, Wee1 becomes inactivated (Fig.5A) and CDK activity jumps to the upper (mitotic) branch of the bifurcation diagram. This analysis suggests that the experiments of Patterson et al. (1) only probed one side of the bistable switch, namely the activation of CDK as the cell both grows and synthesizes additional fusion protein. Since larger cells activate their CDK at smaller fusion-protein level than smaller cells, they observed coexisting low and high CDK activity states at a given fusion-protein level. The alternative steady states of the bistable mitotic switch do not play any role in creating overlapping low and high CDK activity states in Patterson et al.’s experiments. The only necessary requirements to observe overlapping low and high CDK activity states in these experiments are (1) that the fusion-protein threshold for C-CDK activation be size-dependent and (2) that the transition be abrupt. <u>But</u> <u>the switch need not be bistable to reproduce the basic properties of Patterson et al.’s experiments.</u>

**Figure 5:**
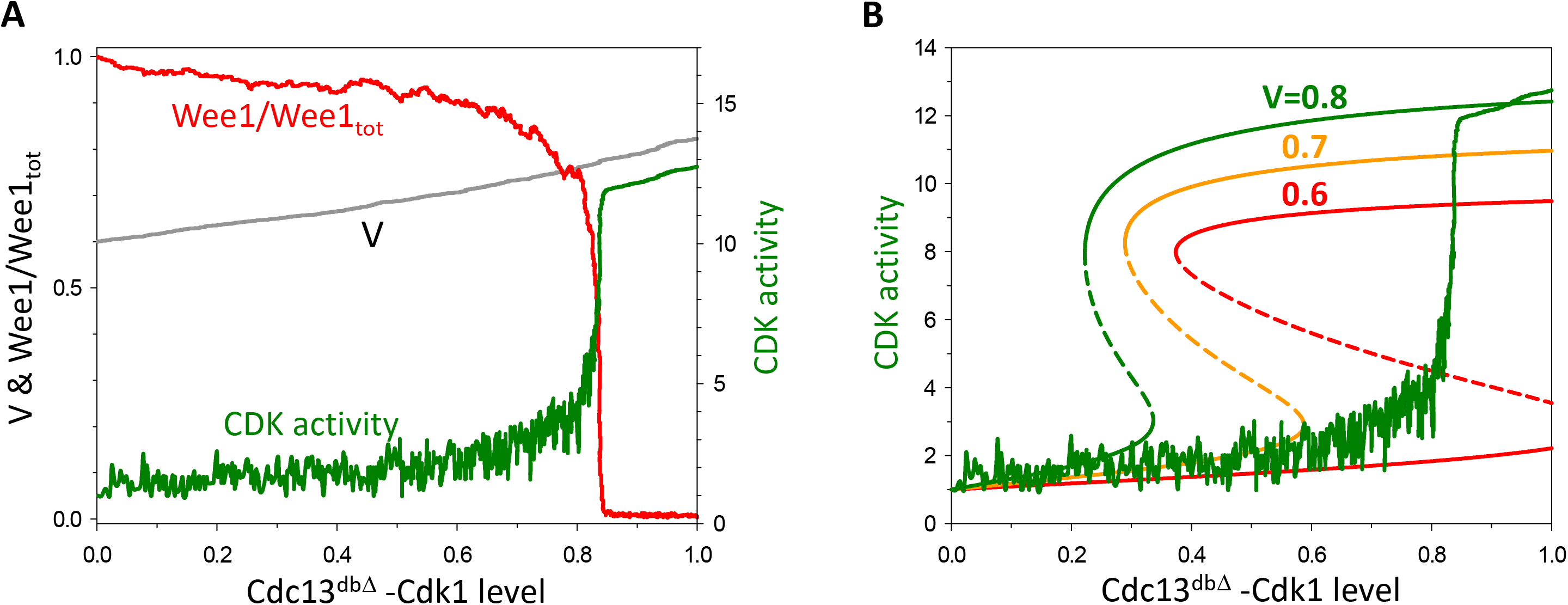
Comparison of the stochastic and deterministic models. (A) Stochastic simulation of mitotic entry induced by fusion protein in WT genetic background for a cell born with volume = 600µm^3^. (B) Overlay of a stochastic simulation on the one-parameter bifurcation diagram.

### Stochastic simulation of a size-controlled reversible switch

To illustrate our argument that a non-bistable switch-like mitotic transition is sufficient to create overlapping interphase and mitotic states, we simulated a model of a reversible switch. Bistability in our model is dependent on a positive feedback loop, whereby C-CDK inactivates Wee1, its inhibitory kinase and stoichiometric inhibitor. To convert our irreversible bistable switch into a reversible sigmoidal switch, we assume that the phosphorylation of the Tyr-modifying enzymes (Wee1 and Cdc25) by C-CDK depends on fusion-protein <u>level</u> rather than CDK <u>activity</u>. Of course, this is a biochemically unrealistic assumption, but it serves to illustrate our point. Under this assumption, the bifurcation diagram (CDK activity vs. C-CDK level) of the mitotic control system becomes sigmoidal in all genetic backgrounds, for both CDK activity (Fig.S4) and for active CDK concentration (Fig.S5). Sigmoidal bifurcation diagrams are characteristic of a reversible, ultrasensitive switch. Observe that the threshold for CDK activation is still cell-size dependent (Fig.S4, S5).

Using this model, we ran stochastic simulations of the induction of non-degradable fusion protein and generated scatter plots of CDK activity vs. C-CDK level, which were sorted into cell-size bins in the same manner as for the bistable model. Stochastic simulations of phosphorylable C-CDK with and without PP2A show wide ranges of overlapping low and high CDK activities at small size (Fig.6A,C)—distributions that are indistinguishable from the bistable model. In addition, both deterministic and stochastic versions of the ‘reversible-switch model’ fit well with the data in Fig.3B of Patterson et al. (1). Stochastic simulations of C-CDK^AF^ with and without PP2A are shown in Suppl. Fig.S6. These simulations prove that a size-controlled, reversible (sigmoidal) switch is sufficient to create overlapping interphase and mitotic states at a fixed fusion-protein level; <u>so the experiments</u> <u>of Patterson et al.</u> (1) <u>do not unequivocally demonstrate that the C-CDK control system is bistable</u>.

**Figure 6:**
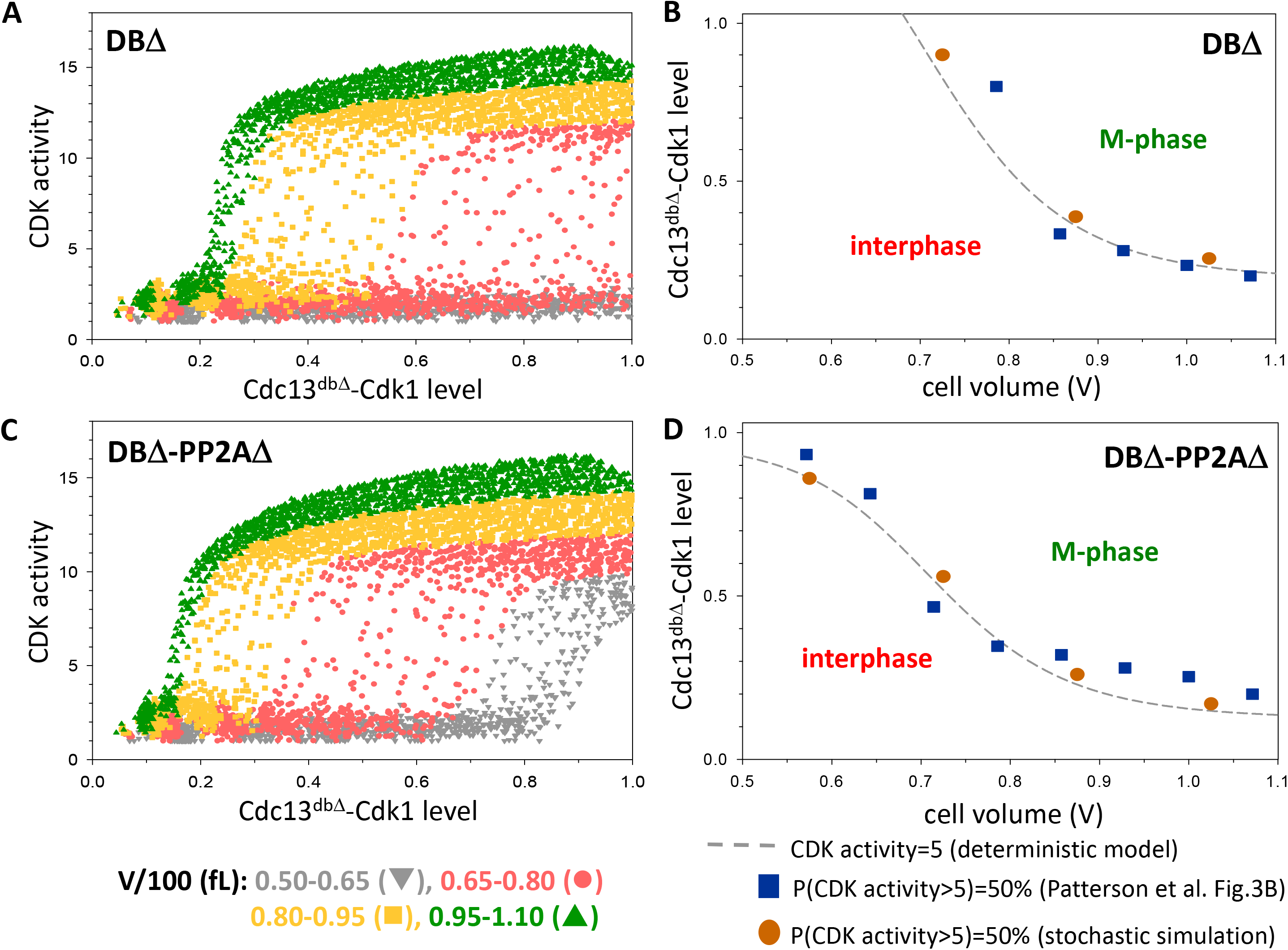
The reversible mitotic-switch model. (A, C) Stochastic simulations of Cdc13^dbΔ^-Cdk1 induction in *pp2a*^+^ and *pp2aΔ*-deleted backgrounds, respectively. (B, D) Cell size dependence of fusion-protein threshold for mitotic entry in *pp2a*^+^ and *pp2aΔ*-deleted backgrounds, respectively. The dashed grey lines depict where CDK activity =5 in the deterministic model. The orange circles and blue squares indicate the fusion-protein levels where more than 50% of cells have CDK activity higher than 5 in our stochastic simulations and in Patterson et al. (1) experiments, respectively.

### Proof of bistability with degradable fusion protein

If the mitotic transition were governed by a reversible switch whose thresholds for CDK activation and inactivation are identical, then the switch would activate and inactivate CDK at the same fusion-protein level regardless of whether the level of fusion protein were increasing or decreasing. To illustrate this point, we performed stochastic simulations of the reversible-switch model starting from the mitotic state (high CDK activity), assuming that the fusion protein is degradable (i.e., the WT strain in Table 1). The results of these stochastic simulations, indicated by the grey data points in Fig.7B-D (overlaid on the previous simulations of mitotic entry induced by non-degradable fusion protein), show that cells inactivate CDK at a level of fusion protein that is only slightly smaller than the level where activation takes place (Fig.7A). This small offset in thresholds for CDK activation and inactivation is caused by time-delays in the activation and inactivation reactions.

**Figure 7:**
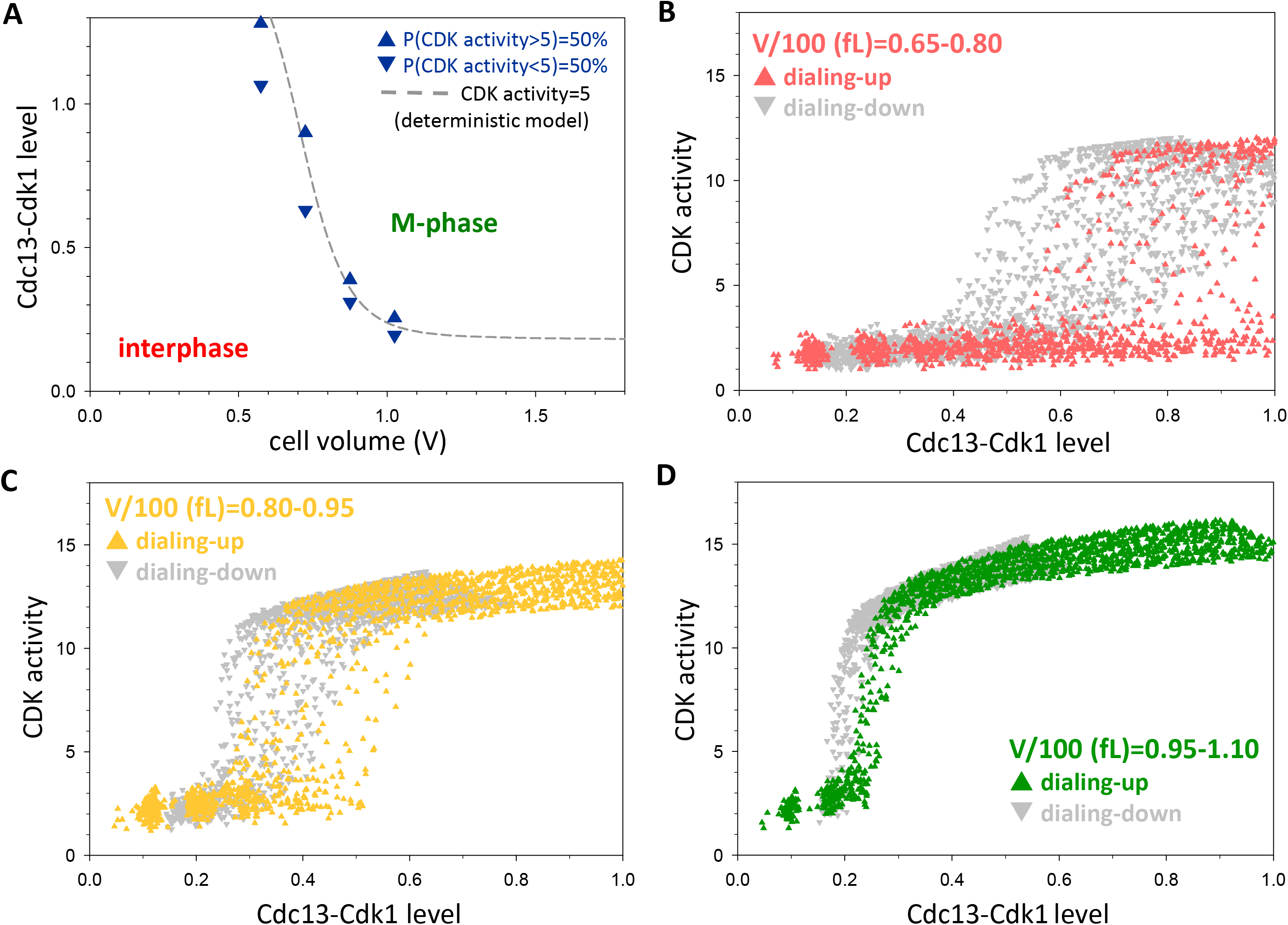
Reversibility of mitotic entry and exit in a reversible mitotic-switch model. (A) The dependence of fusion-protein thresholds on cell size. The dashed line depicts the fusion-protein level where CDK activity =5 in our deterministic model. The triangles (Δ and Δ) indicate the fusion-protein levels where more than 50% of cells have CDK activity higher and lower than 5, respectively, in our stochastic simulations. (B-D) Stochastic simulations of dialing-up the level of Cdc13^dbΔ^-Cdk1 (red, yellow and green data points) and dialing-down the level of degradable Cdc13-Cdk1 fusion-protein (grey data points). Cells are clustered according to size into small (panel B, V=650-800fL), medium (panel C, V=800-950fL) and large (panel D, V=950-1100fL) classes. During dialing-up, non-degradable C-CDK is induced for different times in cells that are initially in the low CDK activity state (interphase); during dialing-down, the level of degradable C-CDK is followed in cells that are initially in the high CDK activity state (M-phase) and are degrading the fusion protein.

**Table 1.** Fission yeast strains used in the experiments of Patterson et al. (1) and discussed in this work.

| Strain notation | Abbrev* | Genotype |
| --- | --- | --- |
| Cdc13<sup>dbΔ</sup>-Cdk1<sup>WT</sup> | DBΔ | *cdc2<sup>ts</sup> TetP:cdc13<sup>dbΔ</sup>-GFP-cdc2<sup>+</sup> ppa2<sup>+</sup> synCut3-mCherry* |
| Cdc13<sup>dbΔ</sup>-Cdk1<sup>AF</sup> | DBΔ-AF | *cdc2<sup>ts</sup> TetP:cdc13<sup>dbΔ</sup>-GFP-cdc2<sup>AF</sup> ppa2<sup>+</sup> synCut3-mCherry* |
| Cdc13<sup>dbΔ</sup>-Cdk1<sup>WT</sup> | DBΔ-PP2AΔ | *cdc2<sup>ts</sup> TetP:cdc13<sup>dbΔ</sup>-GFP-cdc2<sup>+</sup> ppa2Δ synCut3-mCherry* |
| Cdc13<sup>dbΔ</sup>-Cdk1<sup>AF</sup> | DBΔ-AF-PP2AΔ | *cdc2<sup>ts</sup> TetP:cdc13<sup>dbΔ</sup>-GFP-cdc2<sup>AF</sup> ppa2Δ synCut3-mCherry* |
| Cdc13<sup>WT</sup>-Cdk1<sup>WT</sup> | WT | *cdc2<sup>ts</sup> TetP:cdc13<sup>+</sup>-GFP-cdc2<sup>+</sup> ppa2<sup>+</sup> synCut3-mCherry* |

\*We use these abbreviations as shorthand for the four strains studied by Patterson et al. (1) and a fifth strain (WT) proposed by us.

A bistable mitotic switch is distinguished from a reversible switch by a significant difference in cyclin thresholds for CDK activation and inactivation. Repeating the stochastic simulations of the bistable mitotic-switch model with unstable fusion protein (WT strain) starting from the mitotic state (high CDK activity), we obtained the grey data points in Fig.8B-D, overlaid on the previous simulations of mitotic entry induced by non-degradable fusion protein. (Note: we do not expect any significant difference in mitotic-entry dynamics between degradable and non-degradable fusion protein.) These panels illustrate the predicted outcome of the Patterson et al. (1) experiments, if they were done with Cdk1^WT^ fused with degradable Cdc13^WT^. We draw attention to another prediction of the bistable model that the fusion-protein threshold for CDK inactivation is much less size-dependent than the threshold for CDK activation. As a consequence, the hysteresis effect is strongly cell-size dependent and most apparent for small cells which enter into mitosis only at a high level of fusion protein (Fig.8A,B). Based on these results, <u>we propose that a Patterson et al. protocol using fusion</u> <u>protein with an intact cyclin destruction-box could provide unequivocal evidence for bistability of</u> <u>the mitotic switch in fission yeast.</u> Although cyclin degradation happens before cell division takes place, it might be useful to equip the cell with a thiamin-repressible *cdc11* gene (*nmt-cdc11*) and repress Cdc11 expression coincident with the fusion protein induction, which will block subsequent cell septation and separation.

**Figure 8:**
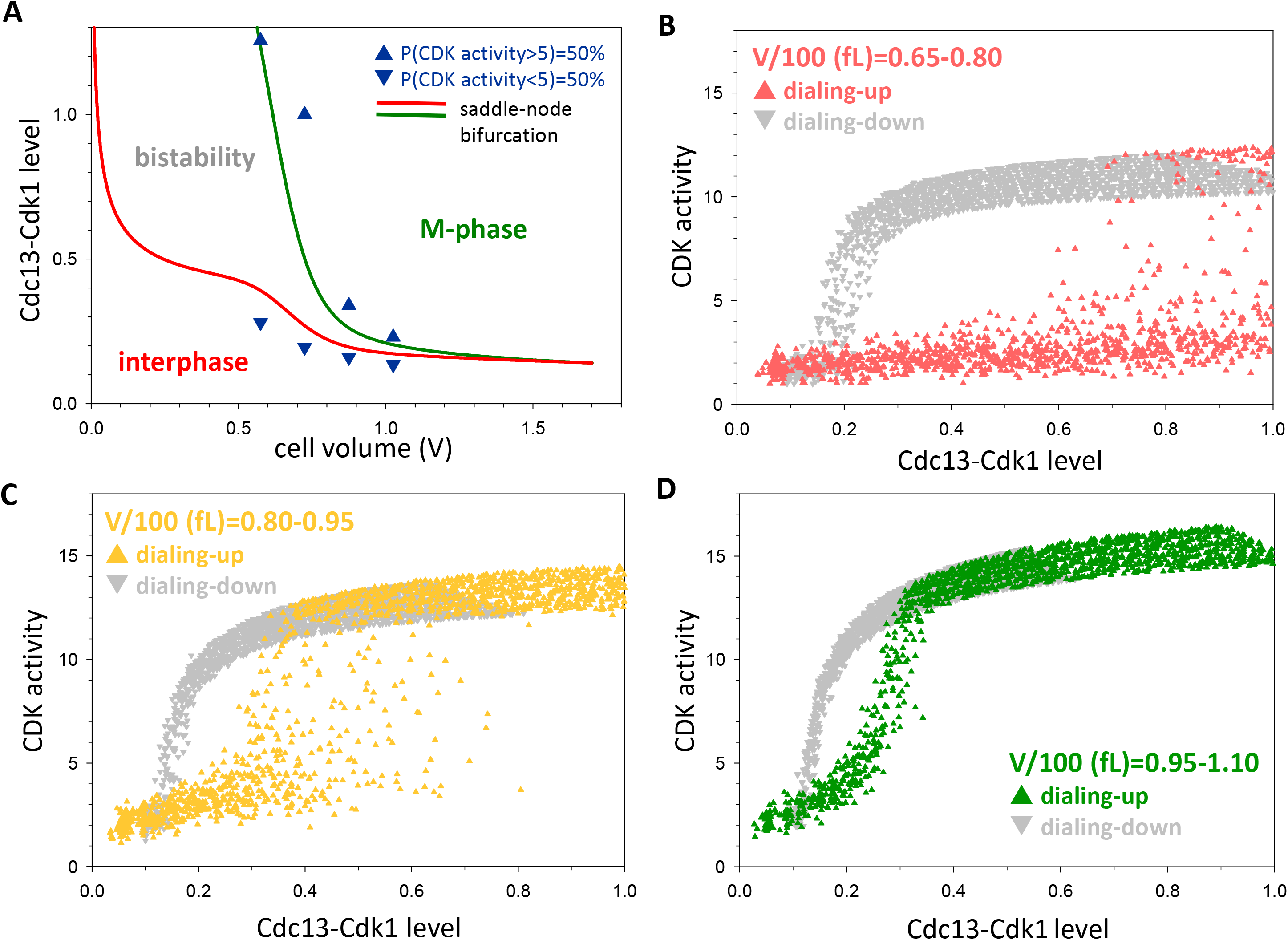
Hysteresis in the bistable mitotic-switch model. (A) The dependence of fusion-protein thresholds on cell size. The green and red lines are the saddle-node bifurcation points in our deterministic model where CDK activity abruptly rises and falls, respectively. The triangles (Δ and Δ) indicate fusion-protein levels where more than 50% of cells have CDK activity higher and lower than 5, respectively, in our stochastic simulations. (B-D) Stochastic simulations of mitotic entry (dialing-up) and exit (dialing-down) with stable (Cdc13^dbΔ^-Cdk1) and unstable (Cdc13-Cdk1) fusion-protein, respectively. Cells are clustered according to their size into small (panel B, V=650-800fL), medium (panel C, V=800-950fL) and large (panel D, V=950-1100fL) classes. During dialing-up, non-degradable C-CDK is induced for different times in cells that are initially in the low CDK activity state (interphase); during dialing-down, the level of degradable C-CDK is followed in cells that are initially in the high CDK activity state (M-phase) and are degrading the fusion protein.

## Conclusions

The eukaryotic cell division cycle (G1-S-G2-M) proceeds, under most circumstances, by a strict alternation of S phase (DNA replication) and mitosis (separation of the sister chromatids to opposite poles of the mitotic spindle). The proper sequence of these events is assured by ‘checkpoints’ at three major ‘decision’ points: the G1/S transition (commitment to DNA replication), the G2/M transition (entry into mitosis), and the metaphase/anaphase transition (exit from mitosis and back to G1). Each of these transitions is governed by a molecular mechanism that exhibits ‘bistable’ dynamics (26), i.e., two stable steady states: a pre-transition state (OFF) and a post-transition state (ON). Checkpoint mechanisms maintain the bistable switch in the OFF state. Once the checkpoint is lifted and the cell passes the transition (flipping the switch ON), it cannot (easily) revert to the OFF state, but rather, the cell returns to the pre-transition state only by passing through all subsequent phases of the cell cycle. In this sense, progression through the eukaryotic cell cycle is a sequence of irreversible transitions governed by bistable switching mechanisms. This understanding of cell cycle dynamics, which is now widely accepted, has been established first by theoretical considerations (3, 11, 20, 27–30) and later by elegant and convincing experimental tests (8, 9, 13, 14, 16). Crucial to all these tests of the theory is the idea that the experimental protocol must flip the switch in two different directions (OFF to ON, and ON to OFF) by perturbations that leave the cell in the same ‘external’ conditions but the switch in either the ON or OFF state. That is the basic signature of bistability: the coexistence of two stable steady states.

Among the experimental investigations of bistability in the cell cycle, a recent paper by Patterson et al. (1) on the G2/M transition in fission yeast seems to confirm the coexistence of ON and OFF states of cyclin-dependent kinase (CDK) activity in cells of similar sizes. Stochastic simulations of a reasonable model of CDK control in the mutant strains used by Patterson et al. quickly confirmed to us that a bistable control system would predict behavior much like that observed by Patterson et al. However, our simulations confirmed our suspicion that their experimental protocol (using non-degradable cyclin, Cdc13^dbΔ^) limited them to observe the OFF to ON transition, only. Although they observed a broad range of cell sizes for which CDK activity could be either high or low, their results could be a consequence of stochastic fluctuations in a model of a reversible sigmoidal switch, provided the switching point is size dependent. We confirmed this suspicion by stochastic simulations of an alternative, sigmoidal switch without bistability. Thus, we conclude that the results of Patterson et al. (1) do not provide convincing evidence of bistable regulation of the G2/M transition in fission yeast.

Furthermore, stochastic simulations with our two models (a bistable switch and a sigmoidal switch) show that a minor change to the genetic strains used by Patterson et al. could distinguish between bistable behavior and sigmoidal switching. The experiments should be repeated with a degradable version of the cyclin component of their fusion-protein construct (i.e., *cdc13^+^-GFP-cdc2^+^*). Some cells must synthesize a large quantity of the fusion protein (as evidenced by GFP fluorescence) to activate CDK activity (OFF to ON) and enter mitosis. These cells will then degrade the fusion protein, and, although the level of the fusion protein will drop considerably, the CDK activity will not flip back to the OFF state until GFP fluorescence drops below a much smaller threshold for CDK inactivation (ON to OFF). A large difference in OFF-to-ON and ON-to-OFF thresholds should be most evident in small cells induced to enter mitosis, and, if observed, it would provide convincing evidence for bistability in the G2/M transition in fission yeast. Furthermore, our model predicts that the OFF-to-ON threshold decreases much more rapidly with cell size than the ON-to-OFF threshold, so the hysteresis loop of the bistable switch shrinks as cells grow.

## Materials and methods

### Computational methods

#### Deterministic model

The time-rates of change of components were described by ordinary differential equations (ODE). At constant level of fusion-protein (CycB_tot_), the ODEs for different forms of CDKs are given by:

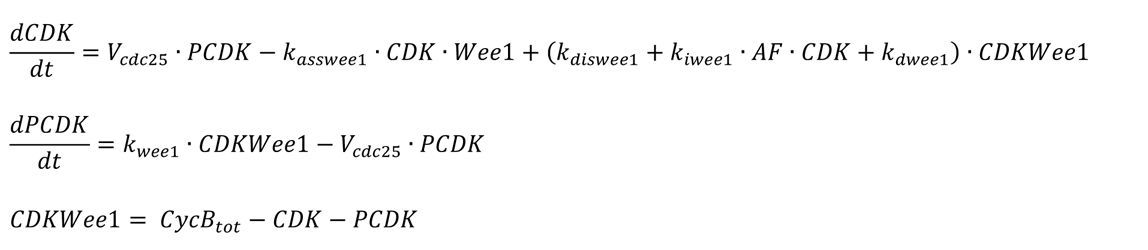

In these equations, CDK and PCDK are the unphosphorylated and phosphorylated forms of the fusion protein. Wee1 forms a complex with CDK (called CDKWee1), which has two fates (see Fig.1): Wee1 phosphorylates CDK and releases PCDK, or a second molecule of CDK phosphorylates the Wee1 subunit, releasing the first CDK moiety + Wee1-P. A third property of the complex CDKWee1 is to function as a stoichiometric inhibitor of CDK activity. The catalytic rate constant, k_wee1_, for the phosphorylation of CDK by Wee1 is set to zero when simulating the Cdk1^AF^ mutant strains.

Concurrently, PCDK is dephosphorylated by Cdc25 at a rate dependent on the phosphorylation state of Cdc25:

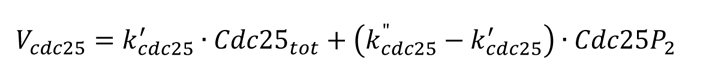

We assume that doubly-phosphorylated Cdc25 has a higher activity than the mono-and un-phosphorylated forms, k_25_” > k_25_’.

The changing concentration of the tyrosine kinases, Wee1 and Mik1 (here treated as a single variable, Wee1) is described by:

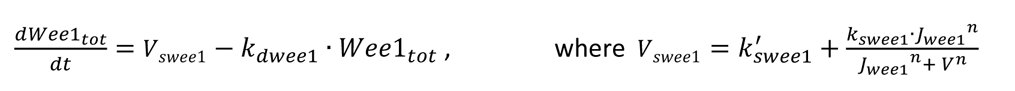

because the synthesis of Wee1 (1^st^ term in V_swee1_) is constant per cell volume (22, 23), while synthesis rate of Mik1 is cell-cycle regulated (24, 25), being restricted to cells with small size (2^nd^ term in V_swee1_).

By contrast, total Cdc25 level increases with cell volume, so we write:

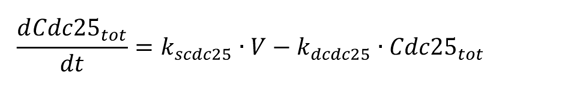

The multisite phosphorylations of Wee1 and Cdc25 are described as distributive and ordered processes with two steps:

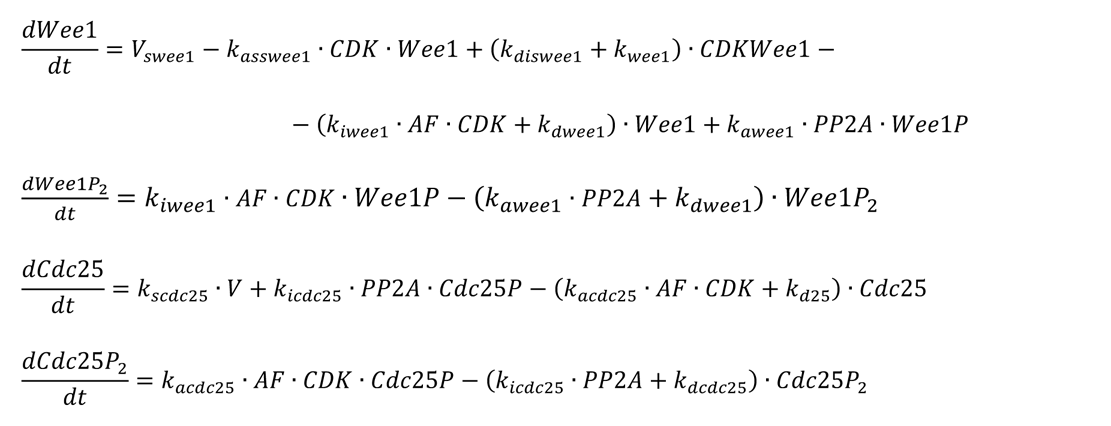

where the mono-phosphorylated forms are calculated by conservation relations:

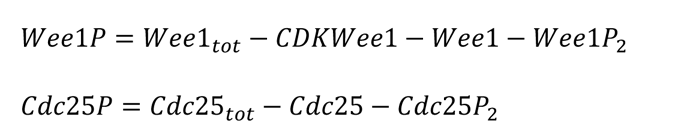

The parameter *AF* defines the specific activity of the unphosphorylated, active fusion protein: *AF* = 1 for the native fusion protein, and *AF* = 0.5 for the AF-mutant, in line with experimental data (31).

#### Calculation of the nucleocytoplasmic ratio of the CDK sensor

To compare our model simulations with the data of Patterson et al. (1), we propose an equation to compute the spatial location of the Cut3-based biosensor (Cut3-mCherry) used to estimate CDK activity in their experiments. Unphosphorylated Cut3 shuttles rapidly between nucleus and cytoplasm, but, when phosphorylated by CDK, Cut3P accumulates in the nucleus. Since CDK is largely nuclear, the sensor (Sand S, for cytoplasmic and nuclear forms of unphosphorylated CDK) becomes phosphorylated in the nucleus, which reduces its rate of export (k_out_’) from the nucleus. The slowly exported, phosphorylated sensor (S_p_) is rapidly dephosphorylated in the cytoplasm. In steady state, the rate of import is equal to the rate of export:

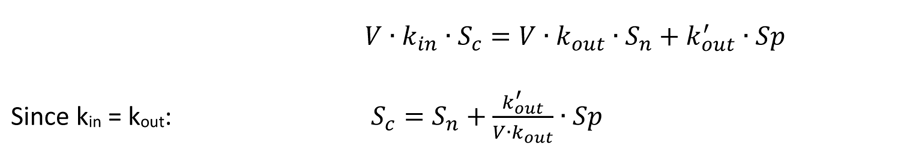

Assuming steady state for the phosphorylated form, the total amount of the sensor in the nucleus is

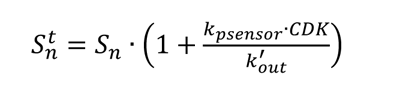

Combining the last two equations, the nucleocytoplasmic ratio of the sensor is calculated by:

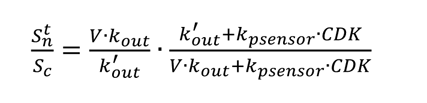

The deterministic model was used to calculate one-and two-parameter bifurcation diagrams using the freely available software XPPAut (32). The values of the parameters are provided in the ‘ode’ file. Rate constants (*k*’s) have a dimension of min^-1^, while other parameters are dimensionless.

### Stochastic model

For time-course simulations, we implemented Gillespie’s Stochastic Simulation Algorithm (SSA) by converting the rates of elementary reactions into propensity functions (17). The model needs to be supplemented with the dynamics of fusion protein synthesis and degradation:

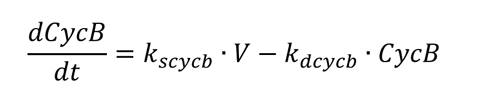

where the overall rate of synthesis is assumed to be proportional to cell volume. Gillespie’s SSA provides exact stochastic simulations for chemical processes governed by mass-action rate laws (reaction rate proportional to the product of the reacting species), as are the reaction rates in our model. Because we do not explicitly model mRNA levels, our protein synthesis terms, like *k*·*V* above, underestimate the fluctuations in protein production attributable to large fluctuations in mRNA numbers (33, 34). This neglect of mRNA fluctuations likely causes the model to underestimate fluctuations in protein numbers.

To convert from ‘concentration units’ in the deterministic model to ‘molecule numbers’ in the stochastic model, we multiply every concentration by volume, for example:

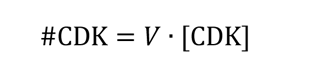

To get a stochastic model with molecular noise comparable to that observed in Patterson et al.’s experiments, we assume that a newborn cell has a volume of ∼500 ‘units’. Since a newborn fission yeast cell is ∼50 μm^3^, 1 ‘volume unit’ is 0.1 μm^3^. To check this assumption, we note that the total ‘concentration’ of C-CDK in large cells (*V* ≈ 100 fL) just entering mitosis is ≈ 0.4 ‘concentration unit’ in the stochastic model (see Fig.2A). This concentration would correspond to #CDK ≈ 400 molecules, which in a cell of volume 100 fL would be a concentration of ∼7 nM. This estimate is much lower than we might expect for CDK concentrations in a mitotic yeast cell (50 nM). In addition to our neglect of mRNA fluctuations, we are also ignoring the contributions of experimental noise to the data; so, it is not unreasonable that we must underestimate the fluctuating numbers of protein molecules in order to get a good fit to experimental observations.

Simulations were run with different initial cell volumes for different lengths of time. For simplicity, we assume that cell volume grows exponentially (doubling time ≅ 140 min) and we set the average cell volume at birth V_B_ = 500 arbitrary units (i.e., 50 fL). Stochastic simulations were started with initial cell volume of 450 (10% below the average birth size) followed by 25 a.u. increments until twice the average birth size (1000 a.u.) is reached. The rate of increase in numbers of C-CDK molecules is set proportional to the actual cell volume. The half-life of C-CDK was chosen ∼6 h, because the fusion protein was made non-degradable by deleting its destruction box. Simulations for each initial cell volume were run for 250 time steps (in increments of 4 molecules of C-CDK synthesized) until 1000 molecules of fusion protein accumulated. Data for cell volume, C-CDK concentrations (number/volume) and nucleocytoplasmic ratio of the sensor were collected at the end of each time step. Each genetic background was analyzed by about 6,000 data points. In summary, stochastic simulations provided a scan of how CDK activity depends on cell size and fusion-protein levels. We also provide the code for stochastic simulations as an ‘ode’ file for XPPAut.

## Acknowledgements

We acknowledge financial support from BBSRC Strategic LoLa grant BB/M00354X/1 to BN.

## Author contributions

Béla Novák: Conceptualization; Software; Formal analysis; Funding acquisition; Writing - original draft, John Tyson: Conceptualization; Formal analysis; Writing - review and editing.

## XPPAut models for fission yeast mitotic switch

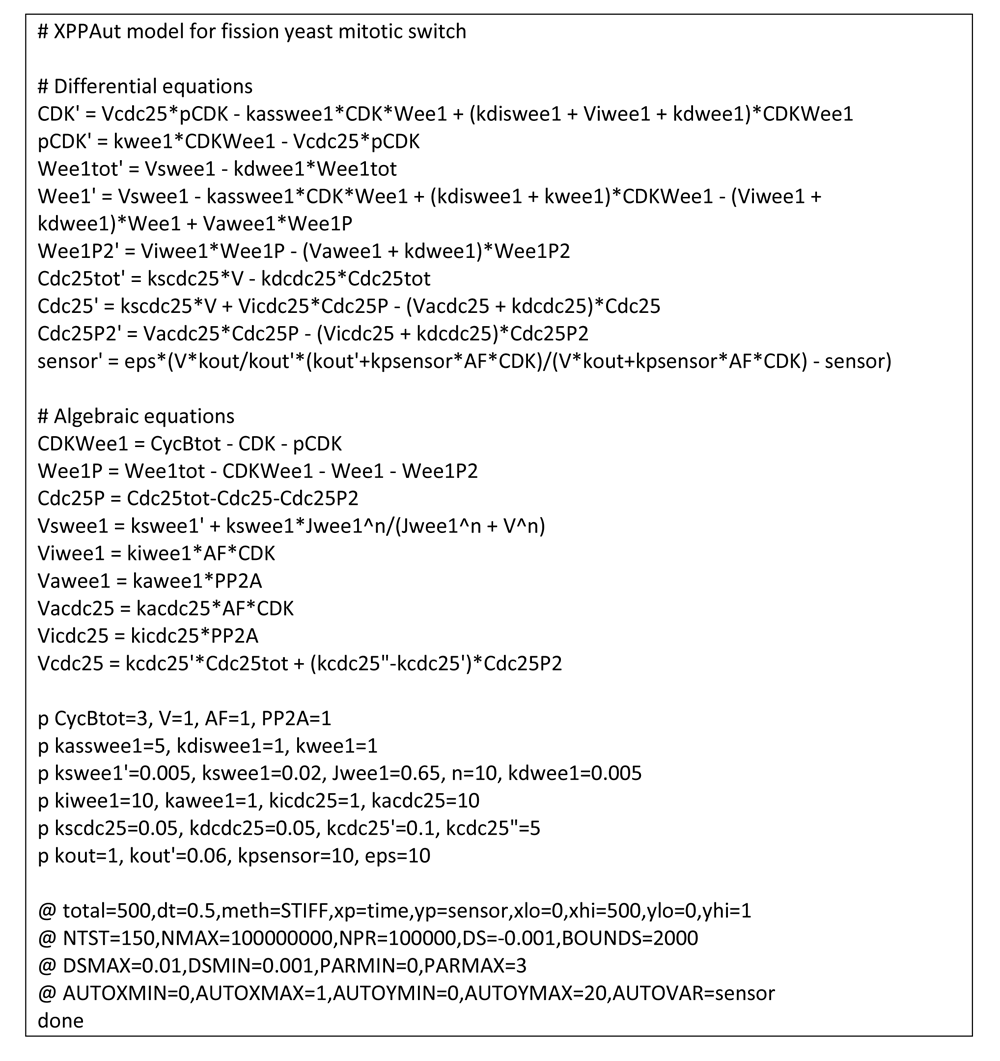
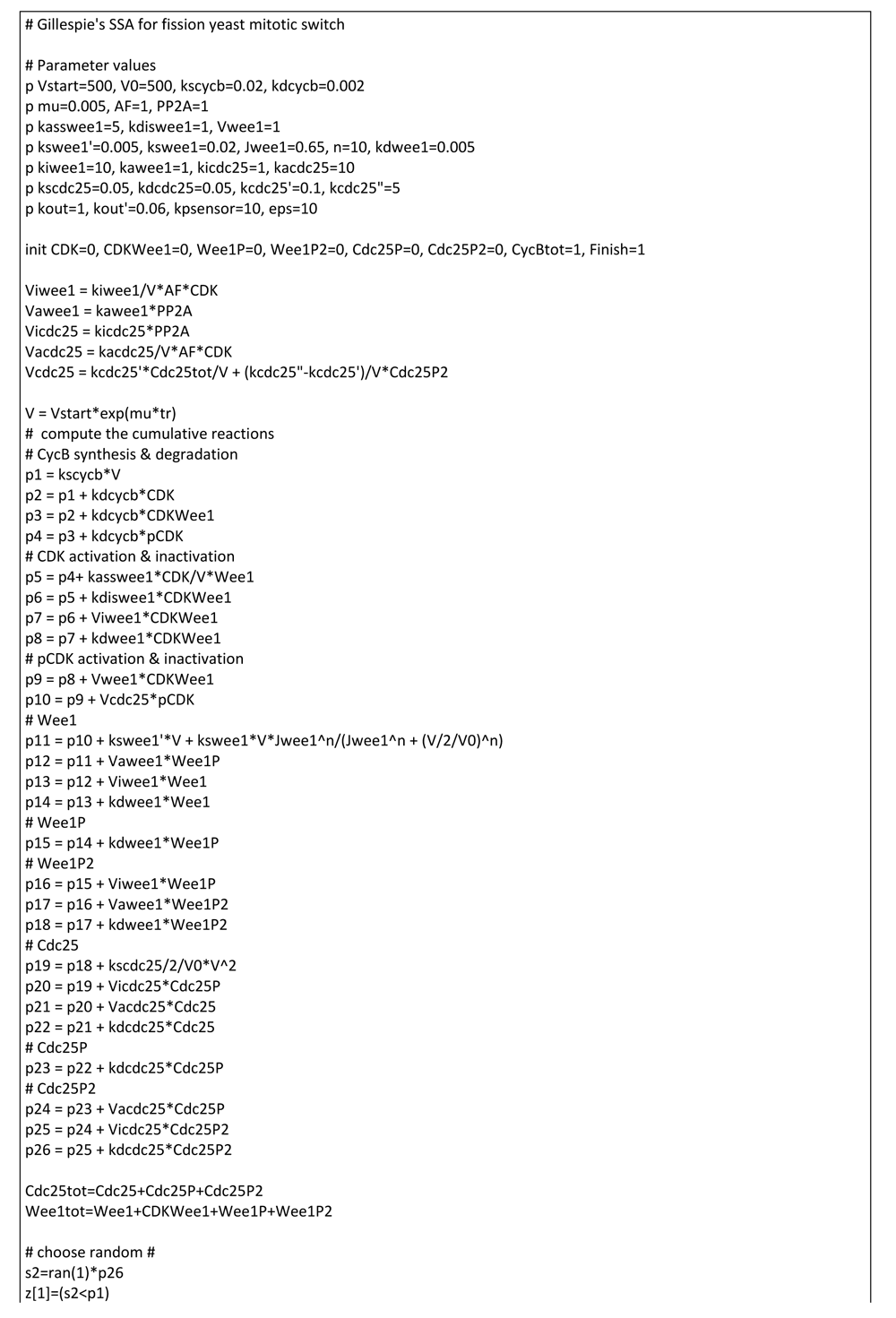
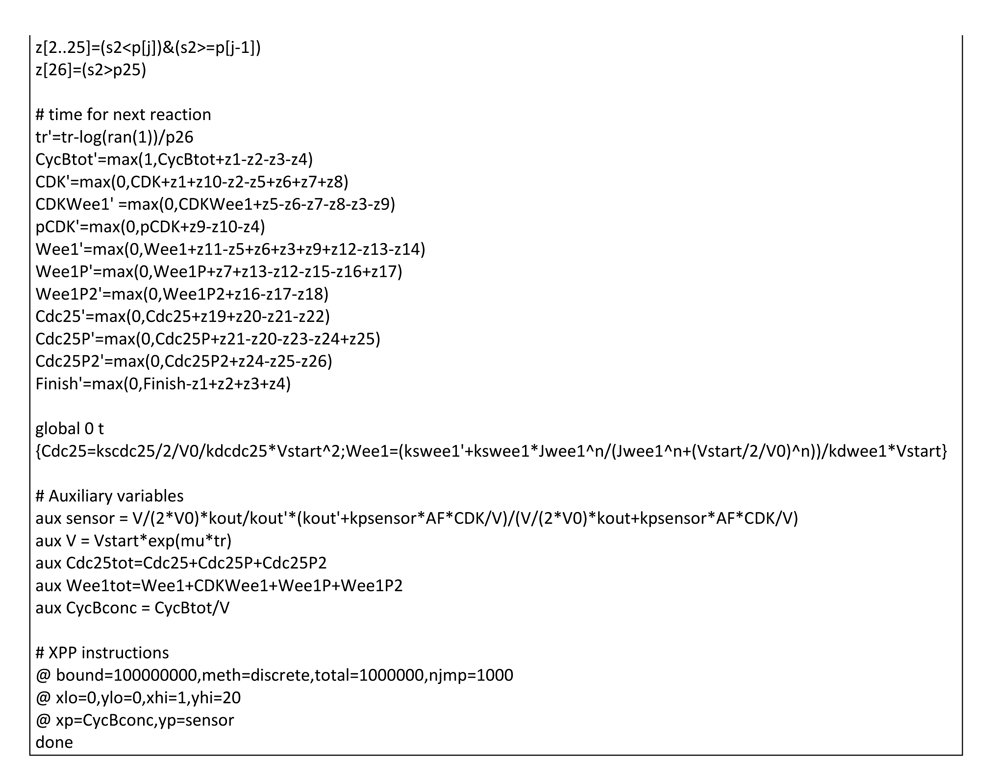

## Supplementary figures

**Figure S1:**
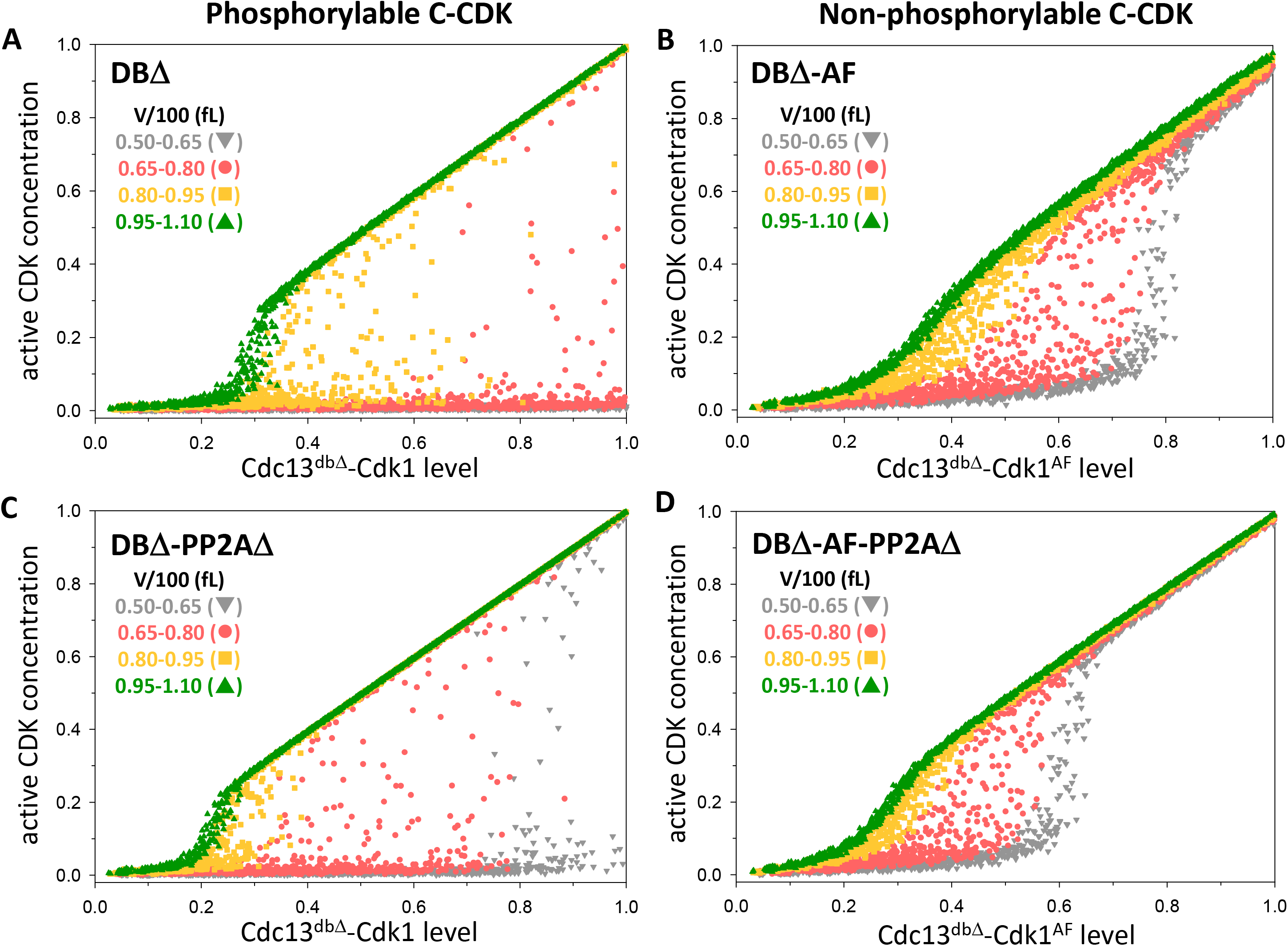
Stochastic simulations of the bistable mitotic-switch model. The level of the active form of CDK is plotted as a function of fusion-protein level after induction of Cdc13^dbΔ^-Cdk1 (left column: A & C) and Cdc13^dbΔ^-Cdk1^AF^ (right column: B & D) in *pp2a*^+^ (top row: A & B) and *pp2aΔ*-deleted background (bottom row: C & D).

**Figure S2:**
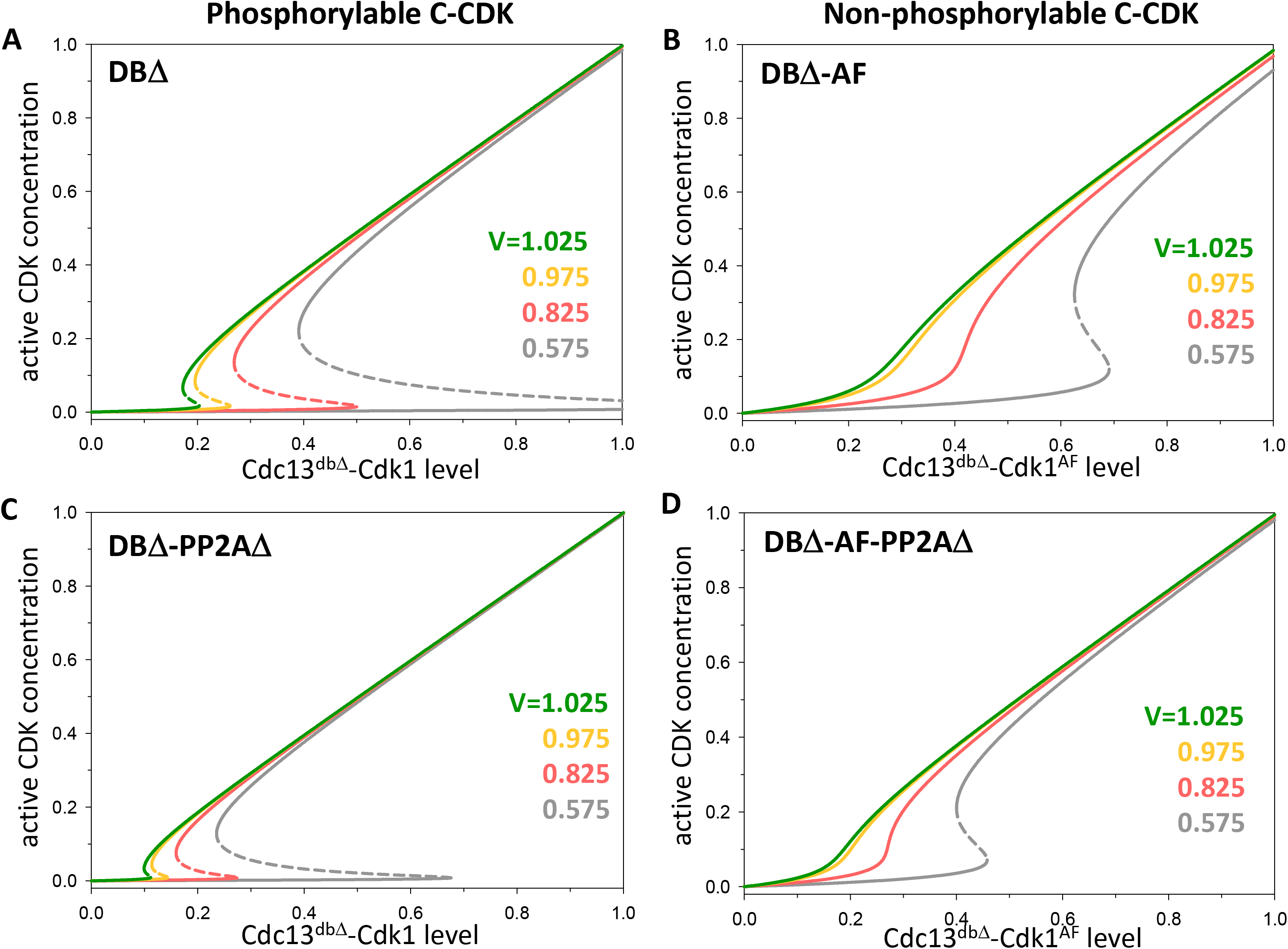
Dose-response curve for dependence of active CDK on fusion-protein level. The level of the active form of CDK is plotted as a function of fusion-protein level after induction of Cdc13^dbΔ^-Cdk1 (left column: A & C) and Cdc13^dbΔ^-Cdk1^AF^ (right column: B & D) in *pp2a*^+^ (top row: A & B) and *pp2aΔ*-deleted background (bottom row: C & D).

**Figure S3:**
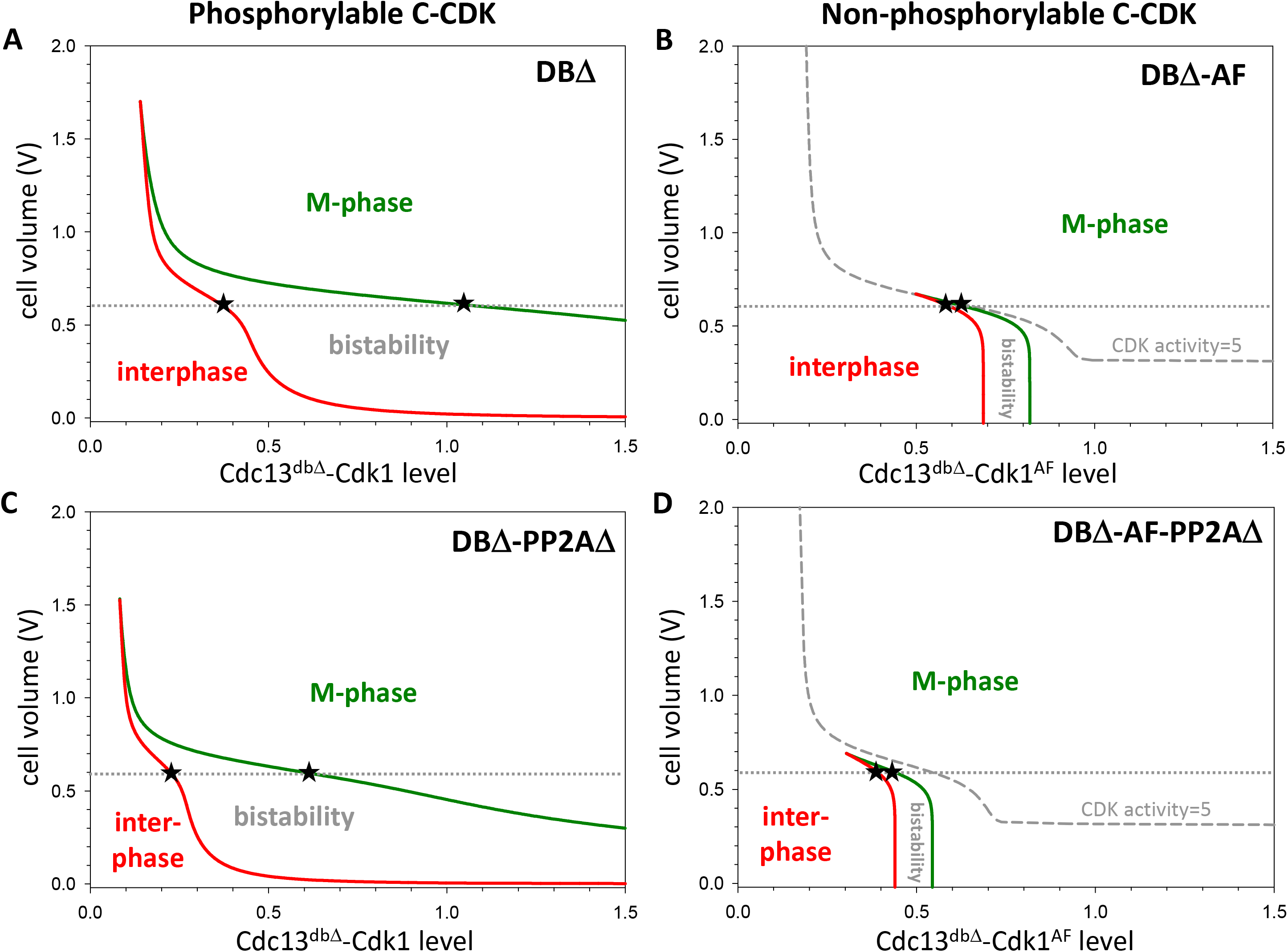
Two-parameter bifurcation diagrams (for the deterministic model) depicting the cell-size dependence of fusion-protein thresholds for mitotic entry (green curves) and exit (red curves. Induction of Cdc13^dbΔ^-Cdk1 (left column: A & C) and Cdc13^dbΔ^-Cdk1^AF^ (right column: B & D) in *pp2a*^+^ (top row: A & B) and *pp2aΔ*-deleted background (bottom row: C & D). The horizontal lines at V=0.6 indicate the width of the bistable domain. The grey dashed lines in panels B and D indicate CDK activity =5 in the Cdk1^AF^ strains.

**Figure S4:**
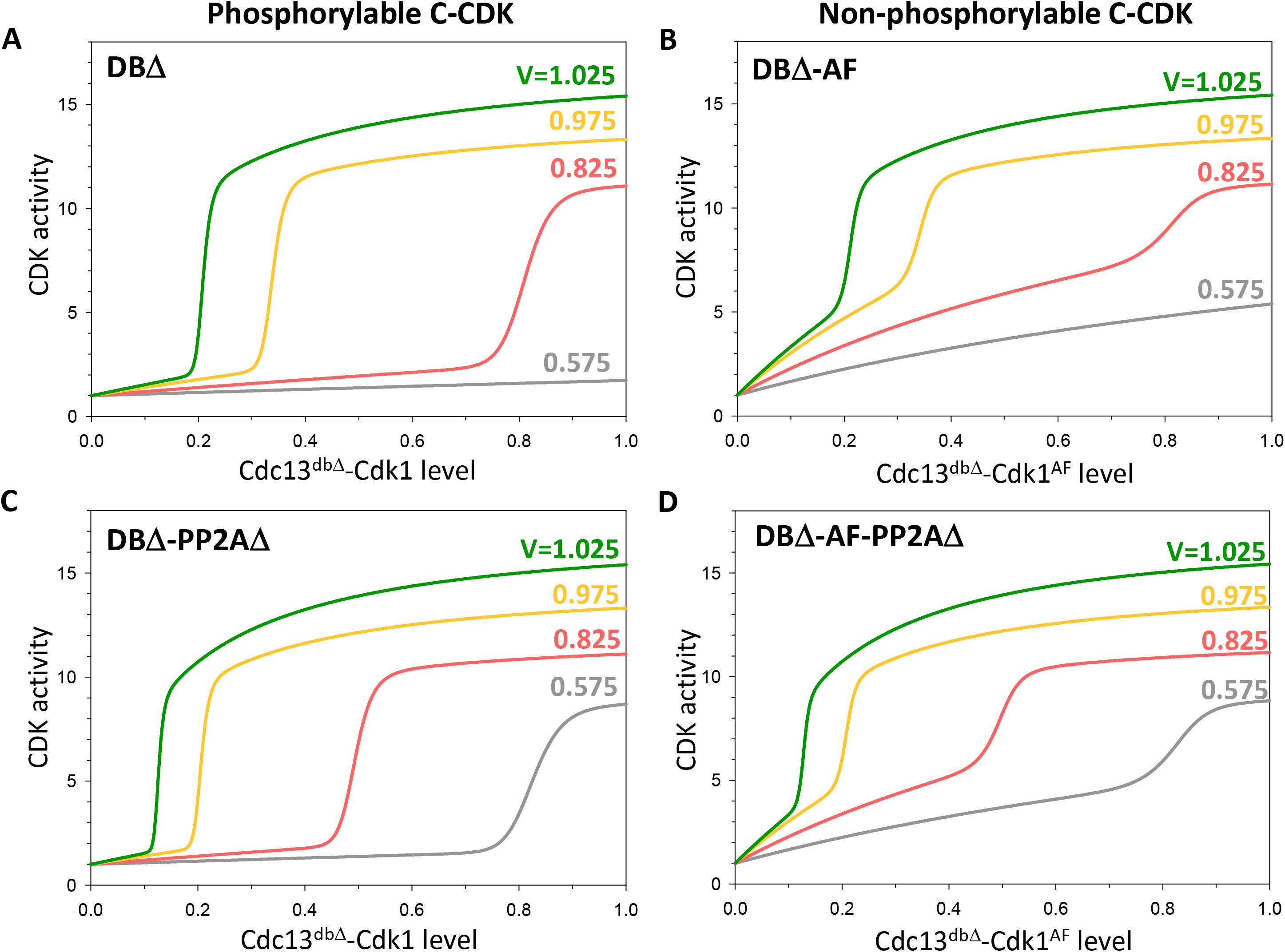
Dose-response curves for dependence of CDK-activity sensor on fusion-protein level predicted by the reversible mitotic-switch model. CDK-activity sensor is plotted as a function of fusion-protein level after induction of Cdc13^dbΔ^-Cdk1 (left column: A & C) and Cdc13^dbΔ^-Cdk1^AF^ (right column: B & D) in *pp2a*^+^ (top row: A & B) and *pp2aΔ*-deleted background (bottom row: C & D). CDK activities are calculated for cell size values in the middle of the cell-size bins.

**Figure S5:**
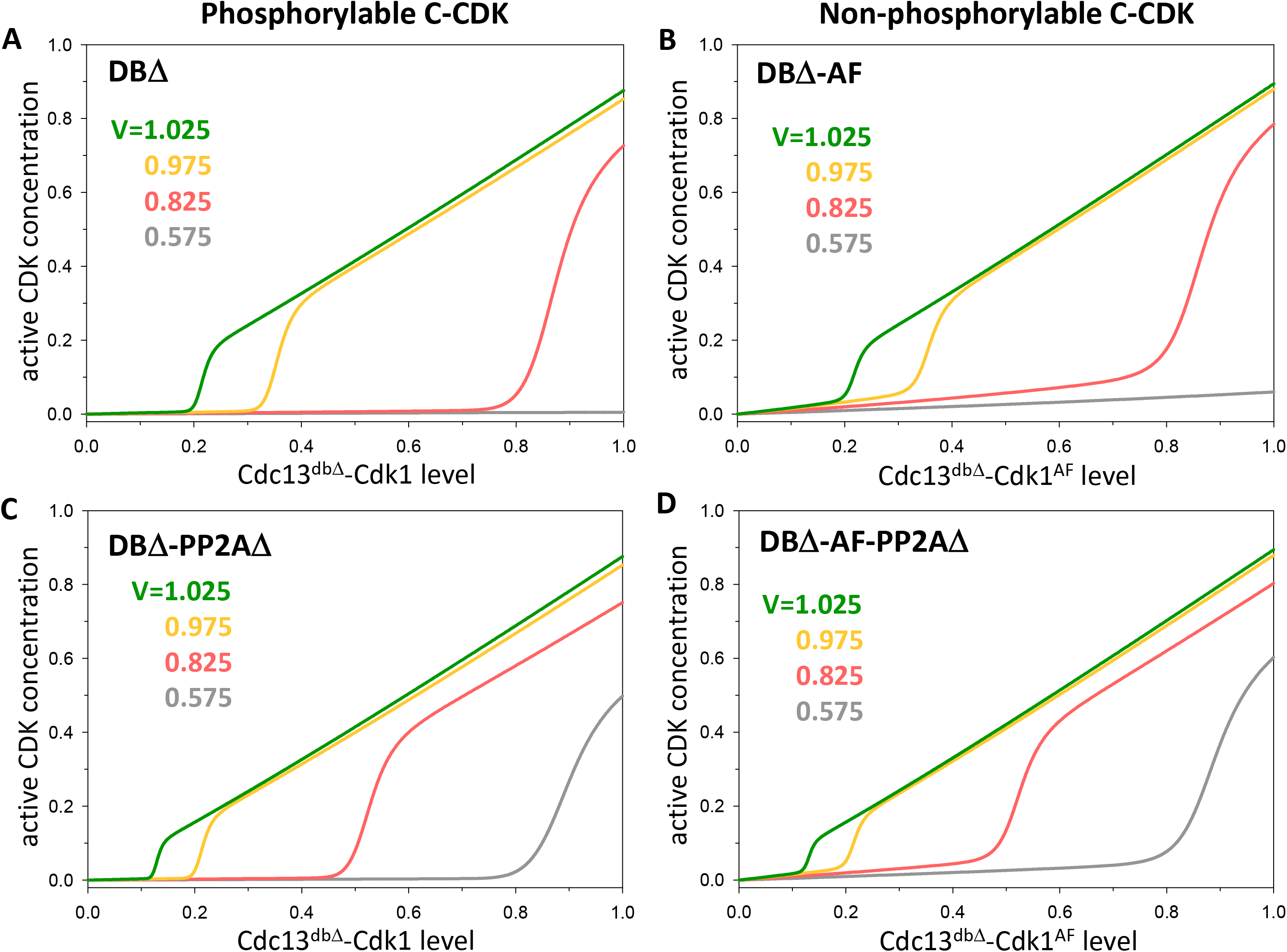
Dose-response curve for dependence of active CDK level on fusion-protein level in the reversible mitotic-switch model. The level of the active form of CDK is plotted as a function of fusion-protein level after induction of Cdc13^dbΔ^-Cdk1 (left column: A & C) and Cdc13^dbΔ^-Cdk1^AF^ (right column: B & D) in *pp2a*^+^ (top row: A & B) and *pp2aΔ*-deleted background (bottom row: C & D). CDK levels are calculated for cell size values in the middle of cell-size bins.

**Figure S6:**
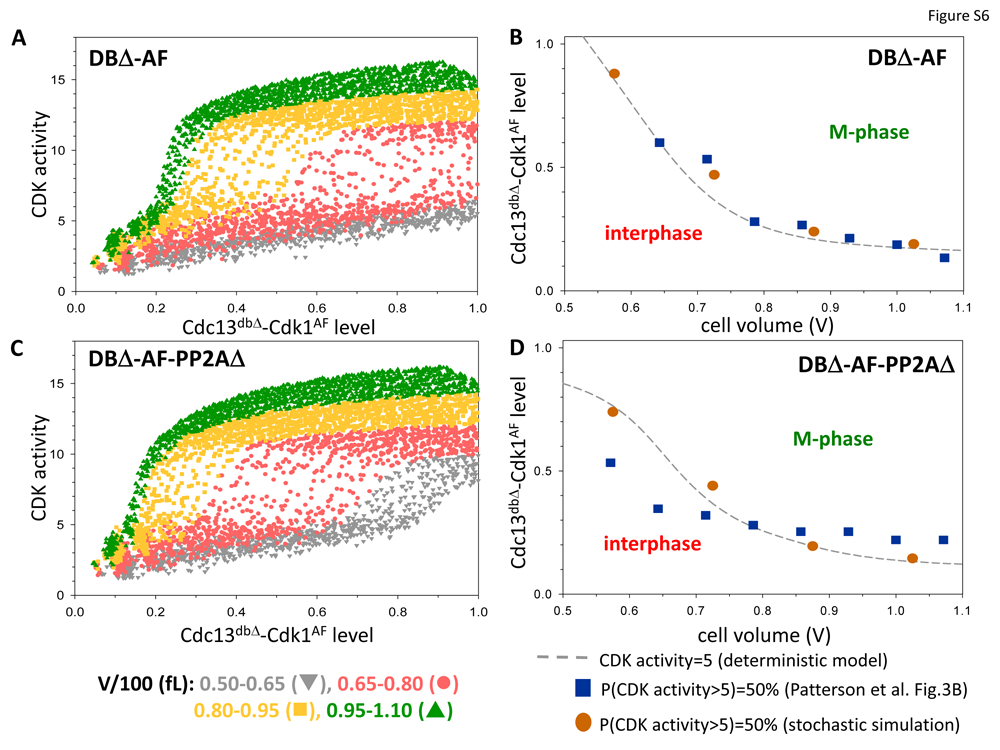
The reversible mitotic-switch model. (A, C) Stochastic simulations of Cdc13^dbΔ^-Cdk1^AF^ induction in *pp2a*^+^ and *pp2aΔ*-deleted backgrounds, respectively. (B, D) Cell size dependence of fusion-protein threshold of mitotic entry in *pp2a*^+^ and *pp2aΔ*-deleted backgrounds, respectively. The dashed grey lines depict where CDK activity =5 in the deterministic model. The orange circles and blue squares indicate the fusion-protein levels where more than 50% of cells have CDK activity higher than 5 in our stochastic simulations and in Patterson et al. (1) experiments, respectively.

## References

1. Patterson JO, Basu S, Rees P, Nurse P. CDK control pathways integrate cell size and ploidy information to control cell division. Elife. 2021;10.

2. Morgan DO. The Cell Cycle: Principles of Control. London: New Science Press; 2007.

3. Novak B, Tyson JJ. Numerical analysis of a comprehensive model of M-phase control in Xenopus oocyte extracts and intact embryos. J Cell Sci. 1993;106 ( Pt 4):1153–68.

4. Novak B, Tyson JJ. Modeling the Cell Division Cycle: M-phase Trigger, Oscillations, and Size Control. J Theor Biol. 1993;165:101–34.

5. Solomon MJ, Glotzer M, Lee TH, Philippe M, Kirschner MW. Cyclin activation of p34cdc2. Cell. 1990;63(5):1013–24.

6. Nurse P. Universal control mechanism regulating onset of M-phase. Nature. 1990;344(6266):503-8.

7. Novak B, Tyson JJ. Qunatitative Analysis of a Molecular Modle of Mitotic Control in Fission Yeast. J Theor Biol. 1995;173:283–305.

8. Sha W, Moore J, Chen K, Lassaletta AD, Yi CS, Tyson JJ, et al. Hysteresis drives cell-cycle transitions in Xenopus laevis egg extracts. Proc Natl Acad Sci U S A. 2003;100(3):975–80.

9. Pomerening JR, Sontag ED, Ferrell JE, Jr. Building a cell cycle oscillator: hysteresis and bistability in the activation of Cdc2. Nat Cell Biol. 2003;5(4):346–51.

10. Tyson JJ, Novak B, Chen K, Val J. Checkpoints in the cell cycle from a modeler’s perspective. Prog Cell Cycle Res. 1995;1:1–8.

11. Chen KC, Csikasz-Nagy A, Gyorffy B, Val J, Novak B, Tyson JJ. Kinetic analysis of a molecular model of the budding yeast cell cycle. Mol Biol Cell. 2000;11(1):369–91.

12. Novak B, Tyson JJ. A model for restriction point control of the mammalian cell cycle. J Theor Biol. 2004;230(4):563–79.

13. Cross FR, Archambault V, Miller M, Klovstad M. Testing a mathematical model of the yeast cell cycle. Mol Biol Cell. 2002;13(1):52–70.

14. Lopez-Aviles S, Kapuy O, Novak B, Uhlmann F. Irreversibility of mitotic exit is the consequence of systems-level feedback. Nature. 2009;459(7246):592-5.

15. Yao G, Lee TJ, Mori S, Nevins JR, You L. A bistable Rb-E2F switch underlies the restriction point. Nat Cell Biol. 2008;10(4):476–82.

16. Rata S, Suarez Peredo Rodriguez MF, Joseph S, Peter N, Echegaray Iturra F, Yang F, et al. Two Interlinked Bistable Switches Govern Mitotic Control in Mammalian Cells. Curr Biol. 2018;28(23):3824–32 e6.

17. Gillespie DT. Stochastic simulation of chemical kinetics. Annu Rev Phys Chem. 2007;58:35–55.

18. Chica N, Rozalen AE, Perez-Hidalgo L, Rubio A, Novak B, Moreno S. Nutritional Control of Cell Size by the Greatwall-Endosulfine-PP2A.B55 Pathway. Curr Biol. 2016;26(3):319-30.

19. Gerard C, Tyson JJ, Coudreuse D, Novak B. Cell cycle control by a minimal Cdk network. PLoS Comput Biol. 2015;11(2):e1004056.

20. Tyson JJ, Csikasz-Nagy A, Novak B. The dynamics of cell cycle regulation. Bioessays. 2002;24(12):1095–109.

21. Novak B, Csikasz-Nagy A, Gyorffy B, Chen K, Tyson JJ. Mathematical model of the fission yeast cell cycle with checkpoint controls at the G1/S, G2/M and metaphase/anaphase transitions. Biophys Chem. 1998;72(1-2):185-200.

22. Curran S, Dey G, Rees P, Nurse P. A quantitative and spatial analysis of cell cycle regulators during the fission yeast cycle. Proc Natl Acad Sci U S A. 2022;119(36):e2206172119.

23. Keifenheim D, Sun XM, D’Souza E, Ohira MJ, Magner M, Mayhew MB, et al. Size-Dependent Expression of the Mitotic Activator Cdc25 Suggests a Mechanism of Size Control in Fission Yeast. Curr Biol. 2017;27(10):1491–7 e4.

24. Ng SS, Anderson M, White S, McInerny CJ. mik1(+) G1-S transcription regulates mitotic entry in fission yeast. FEBS Lett. 2001;503(2-3):131–4.

25. Christensen PU, Bentley NJ, Martinho RG, Nielsen O, Carr AM. Mik1 levels accumulate in S phase and may mediate an intrinsic link between S phase and mitosis. Proc Natl Acad Sci U S A. 2000;97(6):2579–84.

26. Novak B, Tyson JJ. Mechanisms of signalling-memory governing progression through the eukaryotic cell cycle. Curr Opin Cell Biol. 2021;69:7–16.

27. Qu Z, MacLellan WR, Weiss JN. Dynamics of the cell cycle: checkpoints, sizers, and timers. Biophys J. 2003;85(6):3600–11.

28. Qu Z, Weiss JN, MacLellan WR. Regulation of the mammalian cell cycle: a model of the G1-to-S transition. Am J Physiol Cell Physiol. 2003;284(2):C349–64.

29. Thron CD. Bistable biochemical switching and the control of the events of the cell cycle. Oncogene. 1997;15(3):317–25.

30. Thron CD. A model for a bistable biochemical trigger of mitosis. Biophys Chem. 1996;57(2-3):239–51.

31. Rudner AD, Hardwick KG, Murray AW. Cdc28 activates exit from mitosis in budding yeast. J Cell Biol. 2000;149(7):1361–76.

32. Ermentrout B. Simulating, Analyzing, and Animating Dynamical Systems: A Guide to XPPAUT for Researchers and Students. Philadelphia: Society for Industrial and Applied Mathematics; 2002.

33. Pedraza JM, Paulsson J. Effects of molecular memory and bursting on fluctuations in gene expression. Science. 2008;319(5861):339-43.

34. Shahrezaei V, Swain PS. The stochastic nature of biochemical networks. Curr Opin Biotechnol. 2008;19(4):369–74.

